# Social learning of emotion and its implication for memory: An ERP Study

**DOI:** 10.1101/2025.10.02.678576

**Authors:** Sriranjani Manivasagam, Anne Schacht

**Author notes:** Corresponding author: Sriranjani Manivasagam.

## Abstract

Social learning of emotional salience from surrounding social cues is especially advantageous under conditions of uncertainty. Yet, the neural mechanisms underlying this process and its consolidation into long-term memory remain poorly understood. In this two-day EEG study, we examined whether emotional salience from social cues (facial expressions) transfers to perceptually uncertain target images, and whether such learned salience is preserved in memory even after the social cues are removed. On Day 1 (learning session), we found no evidence for automatic emotional salience transfer across trials. Instead, ERP results indicated that social cue use was modulated by participants’ metacognitive state of subjective uncertainty, as reflected in the P1 amplitudes. On Day 2 (test session), recognition memory revealed evidence of additive emotional salience: EPN amplitudes were enhanced for accurately classified positive target images previously paired with social cues. In contrast, LPC amplitudes were reduced for negative target images in the social condition, regardless of classification accuracy. Together, these findings suggest that the influence of social cues is contingent on subjective uncertainty. Social cues enhanced emotional salience when internal valence judgments were strong (as for positive images), but led to increased reliance on the cue—and therefore dampened memory encoding—when internal valence judgments were weaker (as for negative images).

## Introduction

Humans are inherently social beings, thereby making social cues such as facial or vocal expressions, gaze, gestures, postures etc. prevalent in the human environment. Social cues enable social learning through the observation and imitation of others, without the need for direct first-hand experiences ^1–4^. Social learning is assumed to play an indispensable role from early ontogenetic development to the emergence of shared knowledge, ultimately leading to cultural evolution ^5–7^. The foundations of social learning can be traced back to the origins of social cognition, leading to primates’ sophisticated abilities to recognize social cues from conspecifics ^8,9^. Social cues contain a rich variety of information such as a social agent’s object of attention, their intentions and goals ^10^. Recognizing, interpreting and learning from social cues support adaptive behavior for reward acquisition and survival ^11–14^. Social cues trigger enhanced neural processing compared to non-social cues, particularly in brain areas involved in early attention and performance monitoring ^15,16^. Orientation to visual social cues also recruits more cortical structures of the brain, such as the extrastriate, which is key to adapting future responses to environmental objects ^17,18^. Studies on social attention have further pointed out overt shifts of attention towards social cues ^19,10,20–22^. Taken together, the findings indicate that social cues hold a high motivational value, rendering them salient. Particularly, a class of social cues such as facial expressions or vocal sounds (also called affect bursts), conveys rich emotion-related information not only about the social agent’s mental state but also about the value associated with environmental objects (target stimuli). In this study, we examined emotional salience, which arises from social cues such as facial expressions. We further investigated the neural dynamics of social learning of emotional salience using images that depict various environmental objects and scenes, such as flowers, car-crashes, household objects, etc. as target stimuli, and its preservation in memory.

Traditionally, social learning has been suggested to fall within three categories: stimulus enhancement, observational conditioning, and observational learning (imitation and emulation). Consider an example of a person (social agent) reading a book (stimulus) in the garden and smiling (emotionally salient social cue). Stimulus enhancement takes place when an observer’s attention is directed towards the book following the presence of a smile, making the book salient ^23,24^. Observational conditioning occurs when an observer witnesses the co-occurrence of the book and the smile, subsequently getting conditioned to expect a positive outcome from the book ^25–27^. The third category of social learning is observational learning. It involves a change in behavior by imitating the action of the social agent, which in the example above would be to read the same book in the garden. Observational learning also involves emulation, which focuses on the outcome without necessarily copying the actions of the social agent – in this case one could obtain and read the book by visiting a library ^28,29^. A family of mechanisms that subsumes the above categories of social learning has been proposed as associative learning processes ^30–32^. Thus, using associative processes, social learning has been suggested to follow fundamental mechanisms similar to those of non-social learning, with the key difference being the input that drives the learning process, i.e. social cues ^31^. Assuming that associative mechanisms are the underlying principle of social learning, we hypothesize that emotional salience is transferred from social cues to target stimuli, enabling adaptive responses towards the target stimuli.

Previous research has focused on the social appraisal process, in which different social cues influence the evaluation of emotional salience of the target stimuli ^33,34,20,21^. Using the social appraisal conceptual framework, the neural modulations involved in appraising target stimuli based on the emotional salience of social cues have been studied. The appraisal-focused studies allude to expectation-based modulations at the neural level or an instructed perspective-taking strategy, where the participants view the target stimuli from the perspective of the other person (social agent). Although appraisal strategies lead to adaptive responses to the target stimuli, they have been only investigated on a single trial basis. The dynamics and development of the learning of emotional salience over time have been largely neglected ^35–38^. Moreover, the preservation of associated emotional salience in target stimuli in the subsequent absence of social cues have not been investigated. An associative learning framework allows for the investigation of the evolution of emotional salience transfer and its potential long-term maintenance while dissociating the direct effects of the social cues themselves.

Several previous studies have focused on the transfer of emotional salience, often through monetary gains or losses, to neutral stimuli – such as images of scenes, faces, linguistic materials, and abstract symbols that either do not carry inherent emotional salience or are normatively rated to be less emotionally salient. Event-related potential (ERP) evidence indicates that the acquisition of emotional salience associations result in prioritized processing of previously neutral or meaningless stimuli at both early perceptual and subsequent higher-order stages through an assumed ‘value-based attention mechanism’^39^. Further, the effect of transferred emotional salience has been observed during test phases, either on the same day or one day after the learning of associations in behavioral measures as well as in short-^40–43^, mid- and long-latency ERPs ^40,44^. Notably, similar ERP modulations of the transferred emotional salience have been reported for testing with a one-week delay, indicating a persistence of associated emotional salience in long-term memory ^45^.

However, the influence of social cues extends beyond neutral target stimuli to instances, where the target stimuli inherently carry emotional salience, for example due to biological relevance or previous experiences. Therefore, to investigate the transfer of emotional salience from social cues to target stimuli, it is essential to consider the presence of emotional salience from both social cues and the target stimuli. In this study, we examined emotionally salient (happy, sad) and neutral facial expressions as social cues and images that are emotionally salient (positive, negative) and neutral as target stimuli. Neuroscientific research has indicated an identical potency of both emotionally salient facial expressions and images to elicit typical emotion-related ERP markers ^46^. For example, the Early Posterior Negativity (EPN), an ERP component observed at temporal-occipital sites approximately 250 ms after the stimulus onset, has been reported to be augmented in response to both emotionally salient facial expressions and images, presumably reflecting increased selective attention ^46–51^. The EPN is typically followed by the Late Positive Complex (LPC), also named Late Positive Potential, over centro-parietal electrodes, starting around 300 ms after stimulus onset ^52–54,49,55^. This long-lasting ERP response has been assumed to reflect higher-order elaborate and evaluative processes ^46,48^. In addition, some studies have also reported enhanced P1 amplitudes to both emotionally salient facial expressions and images, peaking around 100 ms after stimulus onset and consisting of bilateral occipital positivities. P1 modulations have been functionally linked to the activation of extrastriate visual areas through selective attention at early perceptual stages of stimulus processing ^56,48,57,58^. In this study, we employed ERPs to investigate the neural dynamics of the interaction between the inherent emotional salience of the images and the emotional salience transferred from the facial expressions.

A successful transfer of emotional salience from the social cues to the target stimuli requires two fundamental prerequisites: first, the motivation of the observer to learn from social cues, and second, a certain degree of uncertainty in predicting the emotional salience outcome of the target stimuli. For example, novelty, ambiguity, or perceptual uncertainty (due to less reliable perceptual information) results in a lack of clarity when making judgments about the emotional salience of the target stimuli. Along similar lines, previous studies suggested that social cues are often attended to gain more information about the environment, and the employment of such cues depends on several contextual factors, including prevailing uncertainty, the desire to affiliate with a particular social group, or the judged competitiveness of the social agent providing the cue ^59–61^. Particularly in such contexts, the emotional salience of the target stimuli can be determined by integrating emotionally salient social cues from the environment ^62–64^, leading to behavioral and neural facilitation ^65,66^. Consequently, social learning has been proposed to be a meta-cognitive strategy in humans and other animals ^4,60,67^. Studies have demonstrated that social cues strongly impact early perceptual processing, especially in the presence of the contextual factors mentioned above. Additionally, research on multisensory integration provides evidence that contextual cues are integrated, resulting in prioritized processing at early stages when the target stimuli are presented under perceptual-uncertainty ^66^. Supporting the aforementioned evidence, a study found that social cues can drive selective attention to perceptually uncertain stimuli already at the initial stages of processing ^68^. In addition, previous studies have identified activations in the lower-level processing areas to enable group conformity ^69,70^. A similar observation has been made in an ERP-based study under group pressure, where social cues can strongly impact participants, as evidenced by increased P1 amplitudes in early visual processing when participants aligned with the correct group opinion ^71^. Interestingly, the same study also reported a reduction in P1 amplitudes when participants conformed to an incorrect group opinion, suggesting reduced stimulus processing and discrimination. Based on the above evidence, we expected external motivations in the form of perceptual uncertainty or conformity pressure to result in modulations at the early perceptual stages and higher-order stimulus evaluations.

The present study builds on prior evidence that uncertainty fosters a propensity for social learning. It examines how emotionally salient social cues (facial expressions) influence judgments about the emotional salience of target stimuli (target images). In a learning session, participants performed an explicit judgment of emotional salience of the target images, that is, the participants classified the target images as positive, negative or neutral in a valence classification task under experimentally induced perceptual uncertainty. Perceptual uncertainty was induced by a forward mask that was followed by a brief stimulus presentation for 27 ms. The visual processing of the forward mask competed with the target image and resulted in perceptual uncertainty when judging the valence of the target image (**see Supplementary Information 1**).

We further investigated whether the learned associations, based on valence-congruent combinations of target images and facial expressions during learning, persisted in memory following overnight consolidation, even when the facial expressions were no longer provided in the test session.

The learning session consisted of an explicit valence classification task of briefly presented target images with positive, negative, or neutral valence. The target image was followed by a) happy, sad, or neutral facial expression that was introduced as the emotional responses of previous participants to a certain image (social condition), or b) a scrambled face (control condition). Due to the perceptual uncertainty in the valence classification task, we expected the facial expressions to provide relevant information, resulting in the learning of the target image-facial expression associations. In the test session, an Old/New classification task required decisions on the previously learned target images (old) and distractor images (new), which varied in their emotional salience. The comparison of a) the old target images associated with the facial expressions and b) the old target images associated with the scrambled faces allowed estimating the transfer and preservation of emotional salience in the absence of facial expressions. By monitoring task performance and computing ERPs, we aimed to investigate the behavioral and neural modulation of the hypothesized emotional salience transfer from facial expressions towards the target images and its consequences for subsequent memory.

### Hypotheses (see Table 1. Design Table)

**Table 1.** Design Table with detailed hypothesis, sampling and analysis plan with interpretations for each research question.

| Question | Hypothesis (if applicable) | Sampling plan (e.g., power analysis) | Analysis Plan | Interpretation given to different outcomes |
| --- | --- | --- | --- | --- |
| **Research Question 1.**<br>Does the emotional salience of the facial expressions integrate with that of the target images under perceptual uncertainty during the learning session, and if so, what are the neuro-cognitive mechanisms underlying this emotional salience transfer? | **Null Hypothesis (H0) for Behavioral Performance in the Valence Classification Task and ERP amplitudes:**<br>No difference between experimental conditions (social, condition) as a result of learning. Only effects of target image valence – i.e., emotional compared to neutral images.<br>**H0: Accuracy/ERP amplitudes:** (Emotional images_Social == Emotional images_Control) > (Neutral images_Social == Neutral images_Control)<br>**H0: RTs:** (Emotional images_Social == Emotional images_Control) < (Neutral images_Social == Neutral images_Control)<br>**Hypothesis 1: Valence** | **Sample size:**<br>Maximum sample size of N = 95 based on a power analysis to give a power of 95% to significantly detect the smallest meaningful difference between conditions.<br>**Pre-defined inspection points for interim analysis**<br>N = 60<br>N = 80 | **Hypothesis 1.**<br>Generalized Linear Mixed Model (GLMM) with a binomial error structure and a logit link function with subject, image ID and image ID nested within subject as the grouping factors for the random effects.<br>**Accuracy** ~ *Valence \* Condition \* Trials + Image Complexity + (1 + Valence\*Condition\* Trials + Image Complexity \| subject) + (1 + Condition\* Trials \| image ID) + (1 + Trials \| image in subject)* | **Null Hypothesis –**<br>The emotional salience from the face expression does not transfer to the target images during the course of the task, hence showing no significant differences between the associated conditions (social and control) in both task performance and neural processing. Hence the null hypothesis assumes no effects of emotional salience transfer via associative processes.<br>**Performance measures – Accuracy and RT**<br>During the initial part of the task, we expect better performance (accuracy and RTs) for the social condition irrespective of valence indicating the use of facial expressions in accurate and faster classification of images. However, we expect this effect to evolve across trials into an interaction of valence and condition. We expect that there will be an overall improvement in performance across trials due to perceptual learning and this effect will be greater for emotional |
|  | **Classification Accuracy**<br>*Earlier trials* (Comparable effects across valence in the beginning),<br>**Accuracy:** (Emotional images_Social == Neutral images_Social) > (Emotional images_Control == Neutral images_Control)<br>*Later trials,*<br>**Accuracy:** Emotional images_Social > Emotional images_Control > Neutral images_Social > Neutral images_Control<br>**Hypothesis 2: Valence Classification RTs:**<br>*Earlier trials,*<br>**RTs:** (Emotional images_Social == Neutral images_Social) < (Emotional images_Control == Neutral images_Control)<br>*Later trials,*<br>**RTs:** (Emotional images_Social < Emotional images_Control) < (Neutral images_Social < Neutral | **Stopping point**<br>N = 95<br>An appropriate Type 1 correction will be applied as per the number of peeks into the data (for e.g., Pocock correction).<br>We will also balance the biological sex of recruited participants.<br>**Inclusion criteria:**<br>Age between 18 to 35 years, German or English language proficiency to understand written instructions during the experiment, | **Hypothesis 2. and 3.**<br>Linear Mixed Model (LMM), with subjects, image ID and image ID nested within subject as the grouping factors for the random effects.<br>**Dependent Variable (RT/ERPs: P1, EPN, LPC) ~ Valence \* Condition \* Trials + Image Complexity + (1 + Valence\*Condition\* Trials + Image Complexity \| subject) + (1 + Condition\* Trials \| image ID) + (1 + Trials \| image in subject)**<br>*For the ERP data, after assessing linearity, if the LMMs would not be the best fit, then we will consider non-linear models to fit the data.<br>**Model Fitting**<br>To keep the probability of Type 1 error at the nominal level of .05 and avoid an overconfident model, we will include all theoretically identifiable random slope components and correlations between random intercept and slope into the model. However, we will exclude the correlations from the model if and when they are unidentifiable (recognizable by the majority of the absolute correlation parameters being close to 1). Additionally, if we encounter | images. Further, this difference will be greater for emotional images that are associated with facial expressions due to the additional emotional salience from the facial expressions.<br>**ERP measures – P1, EPN, LPC**<br>We expect the ERP to initially show valence effects which eventually interacts with conditions across trials resulting in enlarged amplitudes for emotional images associated with facial expressions. The interaction of valence and the condition evolving towards the later trials would indicate the transfer of emotional salience from face expression to target images as a result of social learning.<br>**Other Potential Outcome:**<br>If the three-way interaction between valence, condition, and trials does not show significance for both the behavioral performance and the ERP amplitudes, but there is a significant two-way interaction between valence and condition, we will conclude that the transfer of emotional salience across trials has a negligible effect that cannot be detected by the current design. This would also mean that emotional salience transfer from the facial expressions to the target images takes place rather early in the task, which then persists throughout the task, being much greater for emotional images as compared to neutral images (resulting in the |
|  | images_Control)<br>**Hypothesis 3: ERP amplitudes (time locked to images)**<br>*Earlier trials* (Comparable effects across conditions in the beginning),<br>**ERP amplitudes (P1/EPN/LPC):** (Emotional images_Social == Emotional images_Control) > (Neutral images_Social == Neutral images_Control)<br>*Later trials,*<br>**ERP amplitudes (P1/EPN/LPC):** Emotional images_Social > Emotional images_Control > Neutral images_Social > Neutral images_Control | no history of diagnosed neuro-psychiatric disorders and normal or corrected-to-normal vision with lenses or glasses.<br>**Exclusion criteria:**<br>We will exclude participants having less than 30 valid EEG trials per condition in each of the Valence Classification and the Old/New Classification tasks. We will determine the rejection rate (EEG artifact-related dropouts) after collecting data from 20 participants. If more than 6 | convergence issues with our models, we will initially start by changing the optimizers used to fit the model and eventually simplify the models, i.e., in a first step drop the random slope term for the interaction and, if convergence issues persist, drop random slope parameters whose contributions are estimated to be very low.<br>**Inference** - We will use a full versus null model comparison using a likelihood ratio test, where the full model includes both the fixed effects and the random effects and the null model that lacks all the fixed effects of interest, but retaining the control variables and the random effects to be controlled for. For assessing the significance of individual effects in the LMMS, we will use Satterthwaite approximation using the function lmer of the package lmerTest in R and a model fitted with restricted maximum likelihood. In case of GLMMs, we will assess the contribution of individual predictors using the likelihood ratio tests with the drop1 method (test = Chisq). We will obtain 95% confidence intervals of the model estimates by parametric bootstrapping. Additionally, to interpret null findings, we will use a frequentist equivalence testing with upper and | interaction of condition and valence). |
|  |  | participants have to be excluded due to poor data quality, we will pause the data collection and reconsider the experimental design – remove some levels of the factors from the design.<br>We will further exclude trials from behavioral analyses, where no response is given within the predefined time window of 2 seconds, as well as outliers (response time deviating 3 SDs from the mean response time of a given condition per participant).<br>If the criteria<br>for the successful induction of for perceptual uncertainty are not met (Lower accuracy and slower response times in the control compared to the social condition in the first 2 blocks or the performance indicates a ceiling effect for the control condition), we will exclude the participant's dataset from the analysis.<br>Any termination of an experimental session will result in the complete exclusion of that participant's<br>data. | lower bounds as the smallest meaningful effect of 0.2. |  |
| **Research Question 2.**<br>If the expected emotional salience transfer from the facial expressions to the target images occurred during the learning session, would the acquired and integrated emotional salience be preserved in the absence of the facial expressions, and what are the underlying neurocognitive modulations toward recognition memory of the target images in the test session? | **Null Hypothesis (H0) for Behavioral Performance in the Old/New Classification Task and ERP amplitudes:** The recognition memory in task performance and neural difference in terms of ERP between the social and the control condition will not significantly differ, and there will be only effects of emotional compared to the neutral images. In addition, we will see the typical Old/New effects in the respective ERP components (see Hypothesis 7).<br>**H0: Accuracy/ERP amplitudes:** (Old_Emotional images_Social == Old_Emotional images_Control) > Old_Neutral images_Social == Old_Neutral images_Control<br>**H0: RTs:** (Old_Emotional images_Social == Old_Emotional images_Control) < | Sample size, inclusion and exclusion criteria will be the same as that of research question 1, since it is a within-subject design and the same set of participants will perform the tasks on both days. | **Hypothesis 4.**<br>Generalized Linear Mixed Model (GLMM) with a binomial error structure and a logit link function with subject, image ID and image ID nested within subject as the grouping factors for the random effects.<br>**Accuracy** ~ Valence * Condition * Trials + Image Complexity + (1 + Valence*Condition* Trials + Image Complexity \| subject) + (1 + Condition* Trials \| image ID) + (1 + Trials \| image in subject)<br>**Hypothesis 5, 6 and 7.**<br>Linear Mixed Model (LMM), with subjects, image ID and image ID nested within subject as the grouping factors for the random effects.<br>**Dependent Variable (RT/ERPs: P1, EPN, LPC, P300)** ~ Valence * Condition * Trials + Image Complexity + (1 + Valence*Condition* Trials + Image | **Null Hypothesis –**<br>Although the associations on day 1 were learned (reflected by both performance in the learning session, memory test immediately after the learning session and neural measures), the emotional salience from the facial expressions are not preserved in the target images in the absence of the facial expressions, indicating no significant differences between the old target images associated to different conditions (social and control). Hence the null hypothesis assumes no preservation of emotional salience in memory in the absence of the facial expressions during the test session on the following day.<br>**Performance measures – Accuracy and RTs**<br>In the Old/New task, we expect interactions between valence and condition, with greater classification accuracy and faster RTs for the emotional images associated with facial expressions compared to emotional images in the control condition. This would indicate preservation of emotional salience from the facial expressions in the test phase |
|  | (Old_Neutral images_ Social == Old_Neutral images_Control)<br>**Hypothesis 4: Old/New Classification Accuracy**<br>**Accuracy:** Old_Emoional images_Social > Old_Emoional images_Control > Old_Neutral images_ Social > Old_Neutral images_Control<br>**Hypothesis 5: Old/New Classification RTs**<br>**RTs:** Old_Emoional images_Social < Old_Emoional images_Control < Old_Neutral images_ Social < Old_Neutral images_Control<br>**Hypothesis 6: ERP amplitudes (P1, EPN, LPC)**<br>**ERP amplitudes (P1, EPN, LPC):** Old_Emoional images_Social > Old_Emoional images_Control > Old_Neutral images_ Social cue > Old_Neutral images_Control<br>**Hypothesis 7: ERP** |  | *Complexity \| subject) + (1 +Condition\* Trials \| image ID) + (1 + Trials \| image in subject)*<br>*For the ERP data, after assessing linearity, if the LMMs would not be the best fit, then we will use a non-linear model to fit the data.<br>**Model Fitting**<br>To keep the probability of Type 1 error at the nominal level of .05 and avoid an overconfident model, we will include all theoretically identifiable random slope components and correlations between random intercept and slope into the model. However, we will exclude the correlations from the model if and when they are unidentifiable (recognizable by the majority of the absolute correlation parameters being close to 1). Additionally, if we encounter convergence issues with our models, we will initially start by changing the optimizers used to fit the model and eventually simplify the models, i.e., in a first step drop the random slope term for the interaction and, if convergence issues persist, drop random slope parameters whose contributions are estimated to be very low. | which aids recognition memory.<br>**ERP measures – Saliency effects**<br>For the ERPs (P1, EPN, LPC), the interaction of valence and condition would indicate the potency of emotional saliency from both facial expressions and the target images. We expect greater effects for emotional images associated with facial expressions showing combined effects of inherent emotional salient and associated emotional saliency. This neurocognitive modulation towards target images would suggest preservation of emotional saliency in long-term memory even in the absence of facial expressions.<br>**ERP measures – Old/New effects check**<br>We expect the ERPs (P300, LPC) to show Old/New effects, with enhanced positive-going amplitudes for the Old emotional target images (only control condition) compared to new emotional images. This result would be a check for typical Old/New effects (greater positive-going amplitude of the ERPs for the old emotional images compared to the new emotional images), and would indicate that target images were attended to and processed on day 1, considering the task difficulty of the paradigm, where images are viewed under perceptual uncertainty. |
|  | **amplitudes (P300, LPC) – Old/New effects**<br>**ERP amplitudes (P300, LPC):**<br>Old_Emoional<br>images_Control ><br>New_Emoional images ><br>Old_Neutral images_Control<br>> New_Emoional images |  | **<u>Inference</u>** - We will use a full versus null model comparison using a likelihood ratio test, where the full model includes both the fixed effects and the random effects and the null model that lacks all the fixed effects of interest, but retaining the random effects to be controlled for. For assessing the significance of individual effects in the LMMs, we will use Satterthwaite approximation using the function lmer of the package lmerTest in R and a model fitted with restricted maximum likelihood. In case of GLMMs, we will assess the contribution of individual predictors using the likelihood ratio tests with the drop1 method (test = Chisq). We will obtain 95% confidence intervals of the model estimates by parametric bootstrapping. Additionally, to interpret null findings, we will use a frequentist equivalence testing with upper and lower bounds as the smallest meaningful effect of 0.2. |  |

#### Research Question 1

Does the emotional salience of facial expressions integrate with that of target images under perceptual uncertainty during the learning session, and if so, what are the neuro-cognitive mechanisms underlying this emotional salience transfer?

For the learning session, we expected our hypothesized effects (as described below) to emerge through learning across trials. Accordingly, we tested all effects of our experimental manipulation as an interaction with trials. To specify the direction of the interaction, we divide our hypothesis into earlier and later trials, although the statistical models treated trials as a continuous covariate. As we did not expect any differences in the direction of the hypothesized effects, positive and negative valence were grouped together under the category “emotional images” to simplify the effects’ specification. The experimentally manipulated conditions conveying facial expressions were labelled as ‘social’ and those conveying scrambled face as ‘control’. The trial numbers were determined separately for each target image in their order of appearance and not for each target image valence-condition combination, since learning occurred independently for each target image.

##### Valence Classification Task

1. Behavioral Performance Measures

##### Null Hypothesis (H0): Behavioral Performance Measures (Valence Classification Task)

The valence classification accuracy and the response time will not differ significantly between the social and the control condition across trials. However, we anticipated differences in both accuracy and response time for emotional compared to neutral images. That is, greater accuracy and faster response times for the emotional compared to neutral images irrespective of the associated condition (social or control).

##### Hypothesis 1: Valence Classification Accuracy

At the beginning of the experiment, i.e., earlier trials, we expected an overall greater classification accuracy and faster response times in the social condition compared to the control condition, with comparable performance measures for emotional and neutral images, as the uncertainty induced by brief stimulus durations necessitates the use of facial expressions in the valence classification.

Across trials, we expected an enhancement in overall performance as a result of perceptual learning, with greater performance in emotional compared to neutral images because of their inherent emotional salience. Thus, we expected increasing effects of the target image valence to reveal in the behavioral performance measures over the time course of the learning session, resulting in an interaction of the valence of the target image and condition. We expected emotional images in the social condition to show greater performance gains due to the expected integration of emotional salience from the facial expressions with the inherent emotional salience of the emotional images. This will be followed by the emotional images in the control condition due to the presence of inherent emotional salience.

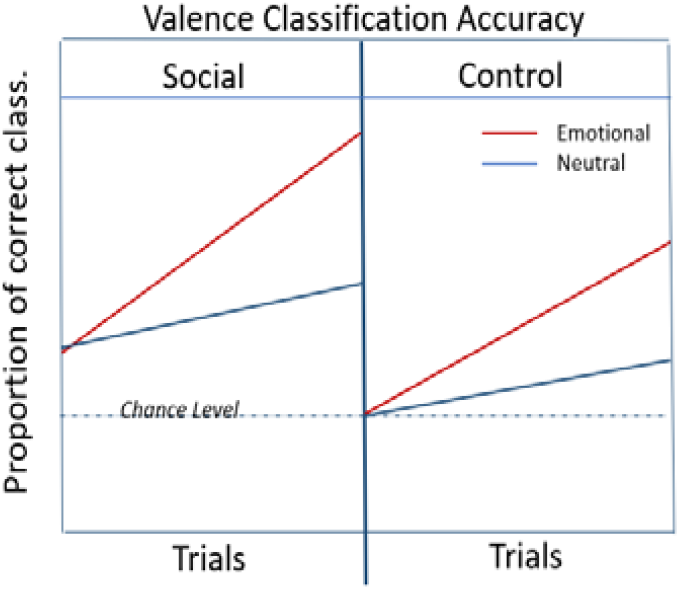

*For earlier trials, we expect:*

(Emotional images_Social == Neutral images_Social) > (Emotional images_Control == Neutral images_Control)

*For later trials, we expect:*

Emotional images_Social > Emotional images_Control > Neutral images_Social > Neutral images_Control

##### Hypothesis 2: Response Times (RTs)

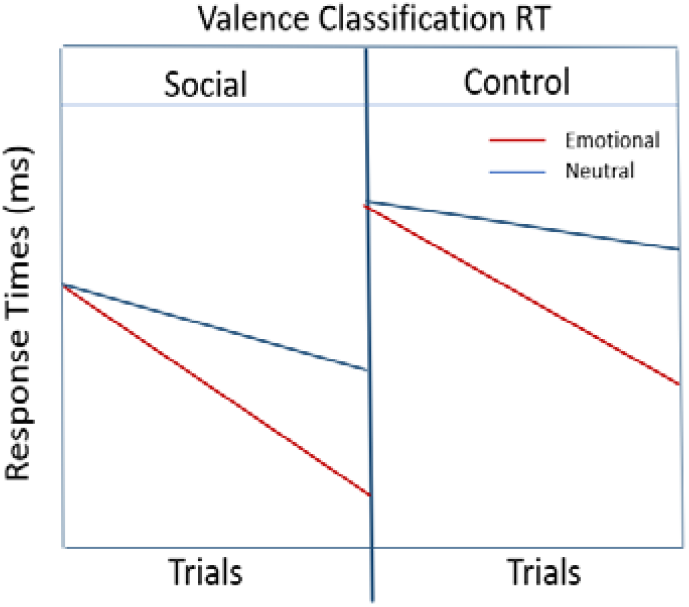

*For earlier trials, we expect:*

(Emotional images_Social == Neutral images_Social) < (Emotional images_Control == Neutral images_Control)

*For later trials, we expect:*

Emotional images_Social < Emotional images_Control < Neutral images_Social < Neutral images_Control

##### Null Hypothesis (H0) : ERP amplitudes for components P1, EPN, LPC

The neural difference in terms of ERP amplitudes will not significantly differ between the social and the control condition. We expected a difference only between the emotional and neutral images. That is, the emotional compared to neutral images will elicit enhanced neural processing, resulting in greater ERP amplitudes irrespective of the associated condition (social or control).

##### Hypothesis 3: ERP amplitudes for components P1, EPN, LPC

At the beginning of the experiment, i.e., earlier trials, we expected comparable ERP amplitudes between the social and control condition since learning of the associations is only expected to occur through the course of the learning session. In contrast to the behavioral performance measures (**see Hypothesis 1-2**), the effect of the associated condition will not be immediately apparent in the ERP components, as they are time-locked to the target image presentation, while the associated condition (social or control) always succeeds the target image within the task.

Thus, across trials during the learning session, we expected effects of condition to unfold over time, resulting in an interaction with the valence of the target images. Specifically, we expected enhanced P1 and EPN amplitudes towards emotional images in the social condition, compared to the control condition. Such results would indicate increased selective attention driven by the combined influence of emotional salience from the facial expressions and the inherent emotional salience of the emotional images. The LPC has been assumed to reflect sustained elaborative stimulus processing and increased memory encoding of emotional stimuli. We thus expected enhanced LPC amplitudes to emotional images associated with facial expressions.

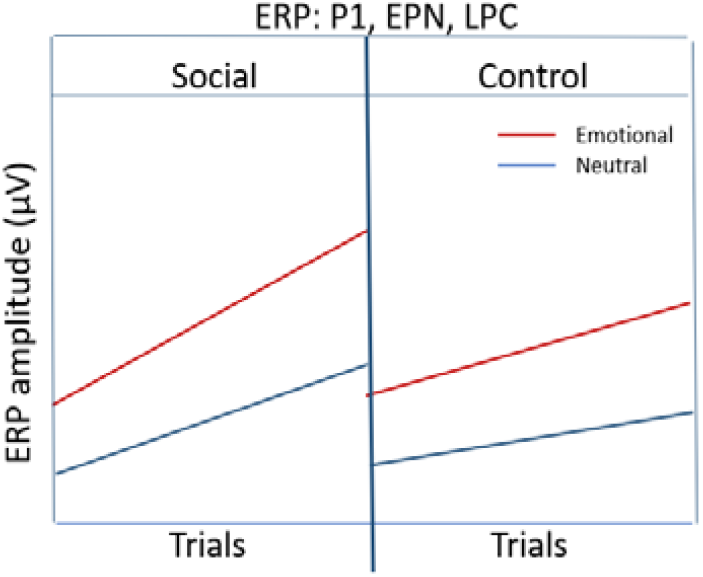

*For earlier trials, we expect:*

(Emotional images_Social == Emotional images_ Control) > (Neutral images_Social == Neutral images_Control)

*For later trials, we expect:*

Emotional images_Social > Emotional images_Control > Neutral images_Social > Neutral images_Control

#### Research Question 2

If the emotional salience transfers from facial expressions to the target images during the learning session, would the acquired and integrated emotional salience be preserved in the absence of the facial expressions, and what underlying neurocognitive mechanisms underlie the recognition memory of the target images during the test session?

Since the new target images only varied in their inherent emotional salience and had no associated emotional salience from the facial expressions, we only compared different conditions of the old target images to assess the preservation of emotional salience through prior associative learning. In addition, we compared the new target images to the old target images in the control condition to verify typical Old/New effects in the ERP components and ensure that the images were attended to. Specifically, we hypothesized that the old target images with emotional salience will elicit more positive-going ERP amplitudes compared to the new target images with emotional salience.

Similar to the learning session, we tested for an interaction of the valence of the target images, condition, and trial in our statistical models. We expected this interaction to capture potential effects of memory extinction over time, if any. However, we only state hypothesis that are relevant with the respect to the research question. Once again, for the hypotheses, the positive and negative valence will be grouped together under the category “emotional images” to simplify the effects’ specification.

##### Old/New Classification Task

1. Behavioral Performance Measures

##### Null Hypothesis (H0): Behavioral Performance Measures (Old/New Classification Task)

The recognition memory and response time between the social and the control condition will not significantly differ and we only expected to see differences between the emotional and neutral images, where the emotional images are remembered better and classified faster compared to the neutral images.

##### Hypothesis 4: Old/New Classification Accuracy

For the Old/New classification accuracy, we hypothesized an interaction between the valence of the target image and condition due to the expected transfer of emotional salience from the facial expressions. This would result in combined effects of emotional salience from the target images and the facial expressions, reflected in better recognition memory for the old emotional images in the social compared to the control condition. Similarly, we expected faster responses to the old emotional images in the social compared to the control condition.

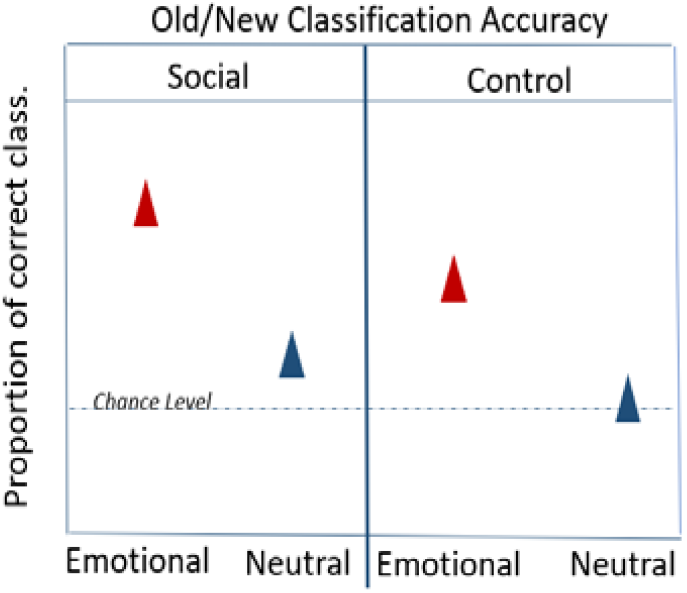

Old_Emotional images_Social > Old_Emotional images_Control > Old_Neutral images_ Social > Old_Neutral images_Control

##### Hypothesis 5: Response Times (RTs)

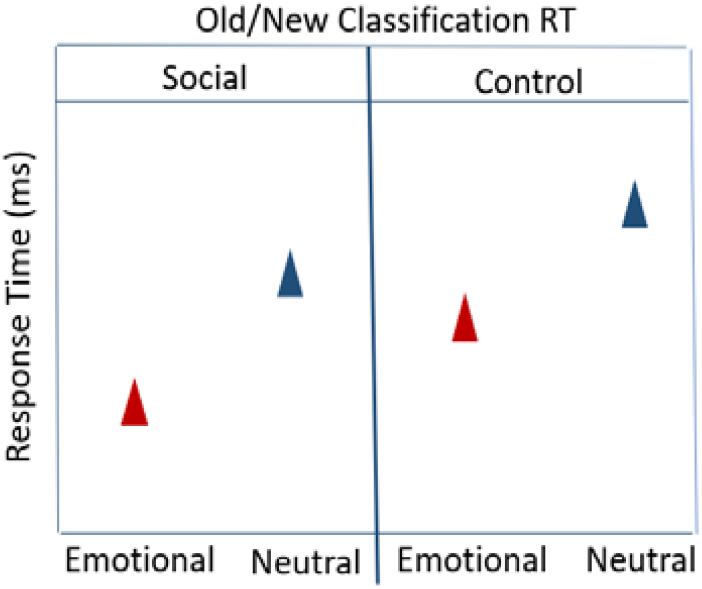

Old_Emotional images_Social < Old_Emotional images_Control < Old_Neutral images_ Social < Old_Neutral images_Control

2. ERP measures

##### Null Hypothesis (H0): ERP amplitudes for components P1, EPN, LPC – Salience effects

The difference in neural activity, indicated by ERP amplitudes, between the social and control conditions will not significantly differ, and we will only see differences between emotional compared to neutral images. Additionally, we expected to observe typical Old/New memory effects in the respective ERP components, with larger positive-going amplitudes for the old emotional images (only the control condition) than new emotional images.

##### Hypothesis 6: ERP amplitudes for components P1, EPN, LPC – Salience effects

We expected an interaction of the valence of the target images and condition, indicating the integrated effects of emotional salience of the facial expressions and the emotional images through prior associations.

In the social condition, old emotional images should gain additional emotional salience due to the integration of the emotional salience from the facial expressions, resulting in an enhanced selective attention and perceptual encoding. This should be reflected in larger P1 and EPN amplitudes for images that had been associated with facial expressions during learning. Similarly, we expected enhanced amplitudes of the LPC amplitudes for emotional old target images associated with facial expressions because they are strongly encoded in memory, leading to better recognition and more allocation of processing resources.

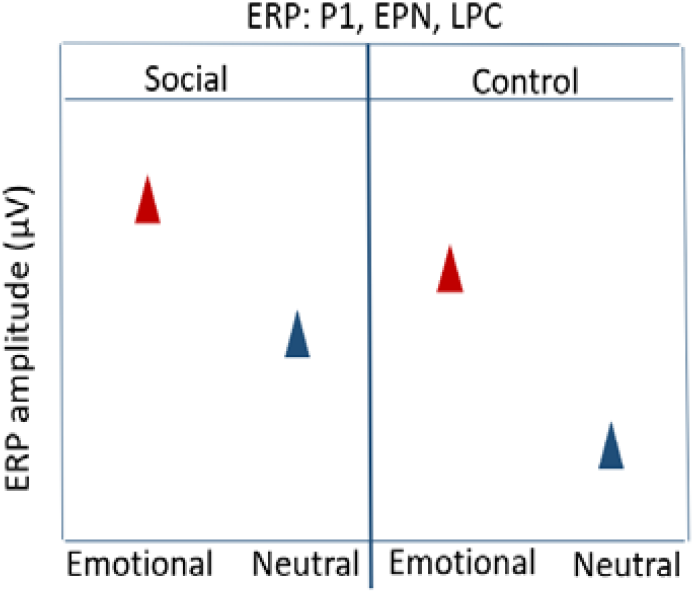

Old_Emotional images_Social > Old_Emotional images_Control > Old_Neutral images_ Social > Old_Neutral images_Control

##### Hypothesis 7: ERP amplitudes for components P300, LPC – Old/New effects

In line with previous studies, we expected typical Old/New ERP effects on the P300 and LPC components. Thus, we expected enhanced positive-going amplitudes to old emotional images (only the control condition) compared to new emotional images.

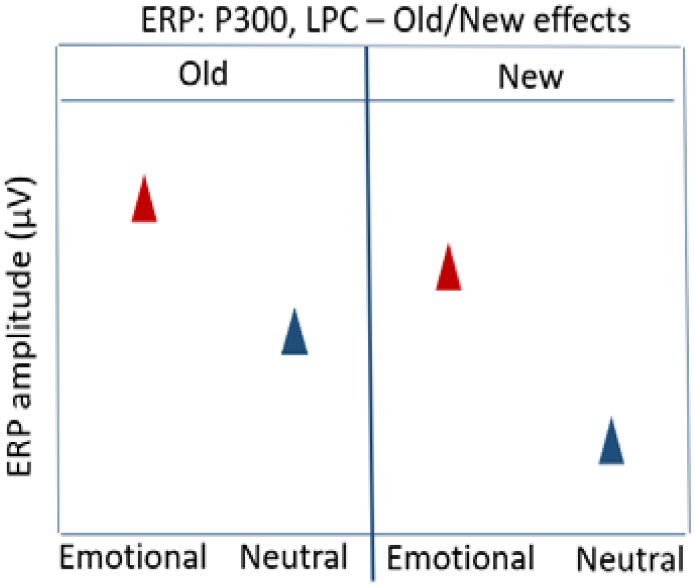

**ERP amplitudes (P300, LPC):** Old_Emotional images_Control > New_Emotional images > Old_Neutral images_Control > New_Neutral image

## Methods

### Ethics information

The study was conducted in accordance with the Declaration of Helsinki, with the approval of the local Ethics committee of the Institute of Psychology at the University of Göttingen. Participants received full information about the procedure and only participated in the experiment after giving their informed consent.

### Design

#### Day1: Part A – Learning session (Valence Classification Task)

The experiment was divided into two sessions conducted on two separate and consecutive days. A self-written script in standard python (v.3.8) utilizing PsychoPy ^72^ was used for stimulus presentation during testing.

The valence classification task consisted of a 2 (social condition/control condition) x 3 (target image valence: positive/negative/neutral) within-subject design. Participants classified the valence of each of the images as either positive, negative, or neutral.

Before the start of the experiment, the participants were prepared for the EEG recording with a 64-electrode EEG cap. The EEG was recorded with 64 active electrodes (AgAgCl) mounted in an electrode cap (Easy Cap™) according to the extended 10-20 system ^73^. For the reference, the common mode sense (CMS) active electrode and for the ground, the driven right leg (DLR) passive electrode was used. The scalp voltage signals were amplified by a BiosemiActiveTwo AD-Box (24 bits; band-pass filter 0.16 - 100 Hz) recorded by the recording software ActiView at a sampling rate of 512 Hz. Additionally, two external electrodes, one on each for the left and right mastoids was used. To record the electrooculography (EOG), the 4 external electrodes that were placed inferior and laterally to the left and right eyes were used.

The participants were seated approximately 55 cm away from the stimulus monitor and instructed to rest their chin on an adjustable chin rest. Electrode offsets were checked to keep them below a threshold of ± 20 mV. Participants then read the instructions at their own pace and completed 10 practice trials. The practice trials contained unique images that did not appear again in the main experiment.

Each trial (see Fig 1a.) started with a centrally presented forward mask for 500 ms with a randomly sampled uniform distribution of jitter ranging between ± 200 ms. The forward mask was centrally presented at a visual angle of approximately 13 x 13 degrees, followed by the target images presented centrally at a visual angle of approximately 2.7 x 2.7 degrees with a 100 x 100 pixel resolution. The target image stimulus was presented for a brief duration of 27 ms. This was followed by a blank screen lasting 973 ms, which allowed for the processing of the briefly presented target image. A fixation cross appeared for 200 ms with a jitter of randomly sampled uniform distribution, ranging between ± 200 ms, to reorient attention to the center of the screen. Following this, the face stimulus (face expression or scrambled face) was presented for 500 ms and a second fixation cross was immediately presented on the disappearance of the face stimulus until the participant’s responded by pressing one of the pre-assigned keys. The faces were presented at a visual angle of about 8 x 9 degrees with a resolution of 300 x 400 pixels. The inter-trial interval lasted for 1000 ms with a jitter of randomly sampled uniform distribution, ranging between ± 200 ms.

**Figure 1.**
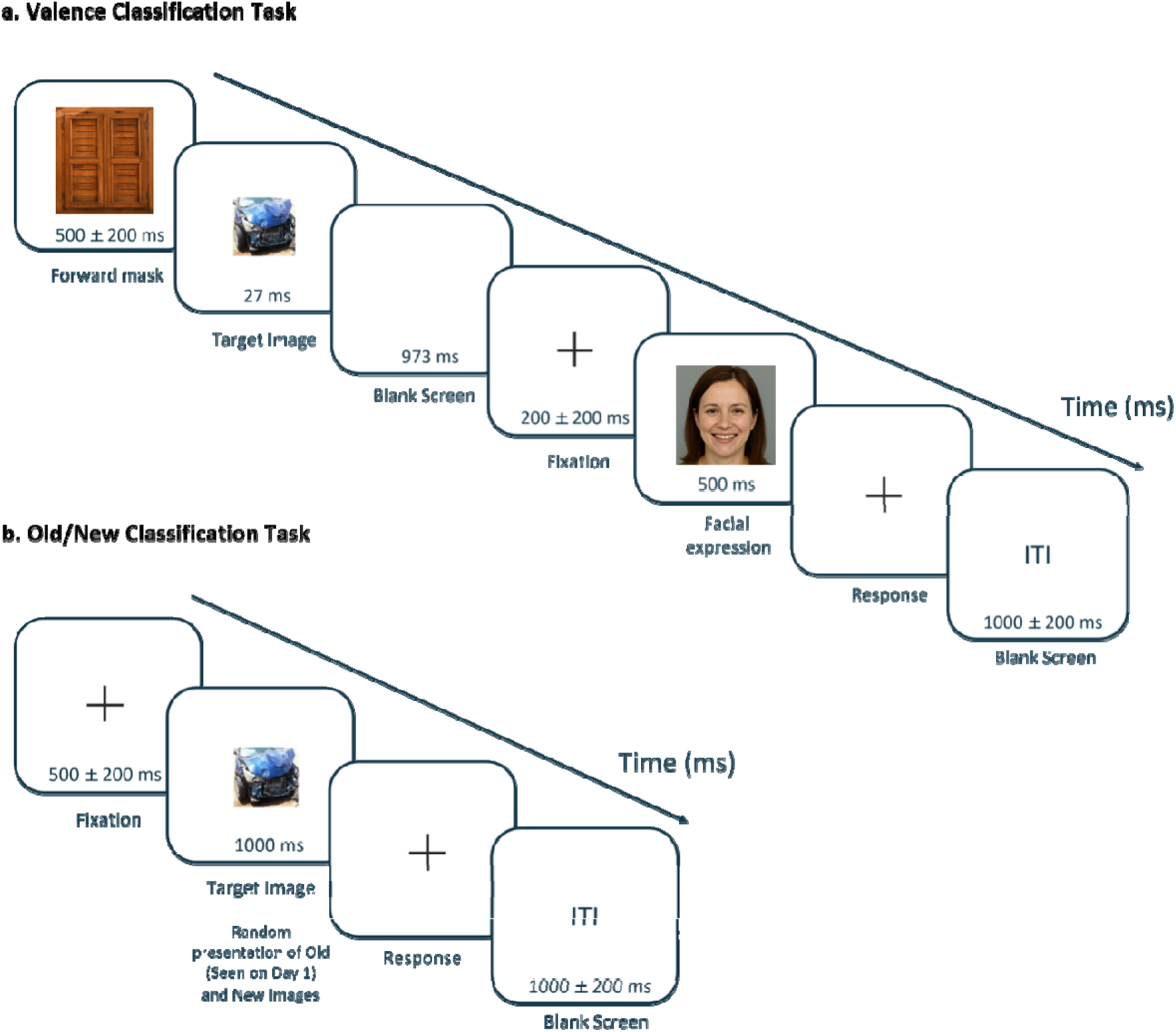
a. Trial scheme of the Valence Classification Task with detailed time sequence. The face shown here is a computer-generated placeholder used for illustration only; actual experimental stimuli were taken from the Radboud Faces Database. b. Trial scheme of the Old/New Classification Task with detailed time sequence.

The valence classification task contained 10 blocks of 42 target images each (14 per valence). The target images were associated with either social cues or scrambled faces with equal probability and the associations remained consistent throughout the task. In the social condition, the target images were always congruently associated to a facial expression (positive image – happy facial expression; negative image – sad facial expression; neutral image – neutral facial expression). Each of the six possible combinations of the target image valence and its associated condition were presented twice within each block. This resulted in 84 trials per block and a total of 140 trials for each of the six combinations across all blocks. There were short breaks between the blocks.

#### Day 1: Part B – Memory Test (Immediately after the Learning session)

Following the learning session, participants performed a memory test which allowed us to examine their acquisition of associations between the target images and the facial expressions. The test consisted of two blocks – the first block assessed recall and the second block assessed recognition memory. The instructions for each of the two blocks of the memory test were presented separately right before the respective blocks. In each block, every target image seen during the learning session were presented once, resulting in 42 trials each. There were 84 trials in total in the memory test.

In the first block (recall), all the target images were presented under perceptual uncertainty (as described in the methods for Day1: Part A – Learning session (Valence Classification Task), without the social or control condition. Participants indicated the correct facial expression that was paired with the target image in the learning session by keypresses. To indicate a scrambled face, participants clicked on the ‘SPACE’ button. The trial scheme for the first block is identical to the Valence Classification Task, except for the second fixation cross and the face.

In the second block (recognition), the target images were presented together with faces (incl. facial expressions and scrambled faces) in identical or different combinations compared to the learning session. Participants indicated if the combination is the same as witnessed during the learning session or not. Half of the target images were presented with the same face as in the learning session and the other half with a different face. The trial scheme for the second block is identical to the Valence Classification Task.

In both blocks, participants were asked to respond as quickly and accurately as possible, within a time-window of 2 seconds.

#### Day 2: Test session (Old/New Classification Task)

The Old/New classification task consisted of a 3 (social condition/control condition/new images) x 3 (target image valence: positive/negative/neutral) within-subject design. Participants were asked to classify the images as old (seen on the previous day) or new images.

Seating of the participant, technical setup, and preparation of the EEG recordings followed the same procedure as in the learning session. Again, participants were allowed to read the instructions at their own pace, followed by 10 practice trials of images that did not appear again in the main experiment.

Each trial **(see Fig 1b.)** began with a centrally presented fixation cross for 500 ms with a jitter of randomly sampled uniform distribution with a range between ± 200 ms, followed by the image for a period of 1000 ms. The target images were presented centrally at a visual angle of approximately 5 x 3 and a resolution of 300 x 200 pixels. After the disappearance of the target image stimuli, another fixation cross was presented until the response of the participant. The inter-trial interval lasted for 1000 ms with a jitter of randomly sampled uniform distribution of a range with ± 200 ms.

There were 10 blocks of Old/New classifications, consisting of 42 old target images presented twice and 42 new target images presented once, resulting in a total of 126 trials per block. This resulted in a total of 140 trials per combination of target image valence and condition for the old set, and 140 trials for each of the target image valence of the new set.

##### Stimuli

Images (**see Table S1. in Supplementary Information 1**) were selected from the IAPS database ^74^. Our selected set followed the general pattern of higher mean arousal ratings for negative images in the database compared to positive ones. The chosen images were further restricted based on their mean complexity scores (as part of the IAPS database). Images with easier figure-ground discriminability (image complexity rating less than 2.5) were selected since ERPs to the early components might be influenced by image complexity ^75^.

The selected images were normalized using an open-source MATLAB code ^76^ to ensure they have the same mean luminance and root-mean-square (RMS) contrast. Facial expressions (happy/sad/neutral) from three male and female portrayers were randomly selected from the Radboud face database ^77^. Note that the face displayed in Figure 1a is a computer-generated placeholder used solely for illustration. The actual experimental stimuli were photographs of actors from the RaFD depicting neutral, happy, and sad expressions.

##### Randomization

Two sets of images, each consisting of 42 images (14 negative, 14 positive and 14 neutral) were defined by random allocation of the images from the pre-selected pool for every participant. One of image sets were used in the Day 1 learning session and associated with either a social or a control condition. For the Day 2 test session, the second image set served as new images and was randomly shuffled along with the old images. For the learning session, every image was randomly assigned to either the social or control condition. The assignments remained constant for a participant throughout the session. Face stimuli was randomly selected from the pre-determined set of face stimuli for each participant, which ensured an equally likely occurrence of every combination of the biological sex of both the participant and the posers in the face stimuli (Female participant – Male face stimuli, Female participant – Female face stimuli, Male participant – Male face stimuli, Male participant – Female face stimuli).

For the second block of the memory test that immediately followed Day 1 learning session, the correct and incorrect associations of the target images-face stimuli was randomly assigned to each target image for each participant.

The assignment of response keys was also counterbalanced for the valence classification, memory test, and the old/new classification. The study did not involve any blinding; hence, data collection and analysis was not conducted blindly to the experiment’s conditions.

### Participants

A total of N = 80 participants were recruited for the study (mean age: 23.33 ± 3.72 SD; 45 female, 34 male, 1 other; 72 right-handed, 6 left-handed, 2 ambidextrous) through a combination of online and offline advertisements, flyers, mailing lists, and social media postings. Participants were compensated at a rate of 8.50 EUR per hour of the experimental session or provided course credits. Participants were native German speakers, with no history of neurological or psychiatric disease, with the ability to read and comprehend instructions in either German or English, no history of diagnosed neuropsychiatric disorders, and normal or corrected-to-normal vision with lenses or glasses.

### Sample size justification

For our sample size calculation, we utilized small effect sizes to simulate artificial data for the present study. As the expected results’ smallest effect sizes pertain to the P1 amplitude, we employed expected P1 amplitude effect sizes to run simulations for the power analysis. We simulated effect sizes for each fixed effect parameter (valence, condition, and trial), as well as their two-way and three-way interactions. For the random effects, we simulated the standard deviations for the random slopes of the fixed effects and their interactions. For the fixed effects, we simulated effect sizes ranging from 0.15 to 0.3, while for the random effects, we simulated standard deviations ranging from 0.3 to 0.6. We opted for smaller effect sizes for the fixed effects and larger values for the random effects because of the relatively new paradigm and the uncertainty in the effect sizes. Large values for the subject-specific random effects were consistent with previous associative learning studies conducted in our lab. We ran 1000 linear mixed models using the simulated effects to assess the probability of significance (p < 0.05) for the expected three-way interaction of valence, condition, and trial for different sample sizes. Based on the simulation conducted, we aimed for a total of N = 95 participants to achieve 95 % power to detect the smallest significant difference between the experimental and control conditions.

#### Inspection of data at pre-defined points

Since the power simulation was based on relatively small effect sizes for the fixed effects, owing the uncertainty of effects in the current study’s paradigm, and relatively large contributions of random intercepts and slopes, we expected the possibility of greater power than revealed by the simulation. Additionally, due to constraints on resources and time, we initially recruited a minimum sample of N = 60, and scrutinized the data for the specified hypotheses. Since the sample size was inadequate, we recruited 20 additional participants, bringing the total to N = 80. Based on the number of data peeks, we employed an appropriate Type 1 correction (like the Pocock correction).

#### Exclusion criteria and data checks

We had planned to recruit additional participants to account for potential exclusions due to too many bad trials/noisy data or potential dropouts between the experimental sessions. Following our lab’s established protocols, we planned to exclude participants with less than 30 valid EEG trials per condition in each of the Valence Classification and Old/New Classification tasks. We planned to determine the rejection rate (EEG artifact-related dropouts) after collecting data from 20 participants. If more than six participants have to be excluded due to poor data quality, we planned to pause data collection and reconsider the experimental design. Depending on the source of the noise (excessive EEG artifacts in specific time windows), we aimed to either stop data collection, or continue after dropping certain levels from the design but keeping the same number of overall trials, which will in-turn increase the trials of the other retained levels.

From both the learning and the test session, trials lacking a response within the predefined 2-second time window, and outliers (i.e., response times that deviate by 3 SDs from the mean response time of a given condition per participant), were excluded from the behavioral analysis.

As a measure of perceptual uncertainty, we had planned to compare the first two blocks of the learning session for differences in valence classification accuracies and response times between the social and control condition for each participant. Lower accuracy and slower response time in the control condition compared to the social condition would have indicated that the experimental design induced perceptual uncertainty. If the above criteria were not met, but the participant’s accuracy for both the conditions was around chance level (33.33 % due to three possible choices: positive/negative/neutral), we had planned to include the participant’s dataset in both the behavioral and the ERP analysis, because although the participant may not have used the facial expressions in their classification, implicit learning of the associations may have taken place. However, if the accuracy levels indicated a ceiling effect, we had planned to conclude that perceptual uncertainty was not successfully induced for the specific participant and exclude the dataset from our analysis. We had planned to determine the exclusion rate after collecting 20 datasets. If more than 1/4 of the participants (5 out of 20) do not meet the above specified criteria and have to be excluded, data collection would have been stopped and our experimental design would have been reconsidered.

For the memory test immediately following the learning session, we evaluated the accuracy and response time of the performance for both the blocks, for each participant. An accuracy above chance level (25% due to four possible choices: happy/sad/neutral/scrambled) in block one and an accuracy above chance level (50% due to two possible choices: same/different) in block two, with a slower response time for the incorrect associations compared to the correct associations was considered as evidence of recall and recognition memory of the associations learned in the learning session. However, if the above criteria are not met, we planned to perform a sensitivity analysis including and excluding the participant’s data for all the proposed hypotheses and check whether the model results change significantly. If there is no significant difference, we planned to pool the participant’s data with the others, since an implicit form of learning may have emerged as preserved memory during the Old/New classification task.

Any termination of an experimental session resulted in the complete exclusion of that participant’s data, and additional participants were recruited to replace the excluded participant’s dataset.

### Analysis Plan

#### EEG preprocessing pipeline

The preprocessing pipeline used followed the standardized procedures used in our lab. The EEG data was preprocessed using the EEGLAB (version 2022.0) ^78^, ERPLAB (version 9.10) ^79^, and BioSig (version 3.8.1) ^80^ toolboxes in MATLAB (R2018a). The signal was referenced to averaged mastoids and was processed through an offline high-pass filter 0.1Hz. Further, a 50 Hz notch filter was also applied to filter out line noise. The data was segmented from -200 to 1000 ms relative to the onset of the target image and corrected to a 200 ms pre-stimulus baseline. Independent Component Analysis (ICA) will be used to remove artifacts such as eye blinks and muscle or channel noise. The data was inspected for any channels with poor signal quality and any removed channels was interpolated using the spherical interpolation algorithm. Then, noisy trials were rejected if voltage values exceed the thresholds of -100 µV or +100 µV within the time window of -200 ms to 700 ms, or if the slope of the voltage trend reaches 100 µV or more within the window of interest for component analysis. Data was defined as statistically improbable and therefore rejected, if a trial contains values exceeding 5 SD of the mean probability distribution (at the channel level as well as for all channels). Finally, the signal was re-referenced to the average of all channels and was used for further statistical analysis.

Based on previous studies, we quantified mean ERP amplitudes at the following time windows (in milliseconds (ms)) and regions of interests (ROIs):

***P1*:** 80-120 ms; occipital cluster: O1, O2, Oz

***EPN*:** 250-300 ms; occipital-temporal cluster: O1, O2, P7, P8, P9, P10, PO7, PO8, PO3, PO4

***LPC:*** 400 - 600 ms; centro-parietal cluster: CP1, CPz, CP2, P3, Pz, P4, PO3, POz, PO4

***P300:*** 300-400 ms; parietal cluster: Pz, P1*, P2*, PO3, PO4

*\*In the stage 1 Registered Report the regions of interest were incorrectly entered as PO1 and PO2*

Given the considerable task difficulty of our paradigm, we modified the time-windows and ROIs after visualizing the actual scalp distribution and latency of the ERP of interest (reported in the results section).

#### Statistical analysis

Statistical analysis was performed using R statistical software (4.2.0 or higher, R Core Team, Vienna, Austria) and RStudio (version 2022.02.2, RStudio, Boston, MA, USA).

To investigate the probability of correctly classifying an image into the predefined valence category and Old/New categories, we used a Generalized Linear Mixed Model (GLMM) with a binomial error structure and a logit link function with subject, image ID, and image ID nested within subject as grouping factors for the random effects. For the response times and ERPs (P1, EPN, P300 and LPC amplitudes) measured in both tasks, we used Linear Mixed Models (LMMs) with subject, image ID, and image ID nested within subject as the grouping factors for the random effects.

##### Fixed and Random effects

All the models used to test the specified hypotheses had an interaction of valence (positive/negative/neutral), condition (social /control or old/new - for the Old/New effects), and trials. Trials were included as a covariate in the fixed effect parts of the model as an interaction with the other factors to assess the effects of learning in the learning session and extinction in the test session. In addition, to control for the effect of image complexity, the complexity ratings obtained from the IAPS database were added to the fixed effects part of model. All the models had ‘Subject ID’, ‘Image ID’ and 22 Image nested within Subject (‘Images in Subjects’) as grouping factors for the random effects. The ‘Images in Subjects’ grouping factor was created by combining the ‘Subject ID’ and the ‘Image ID’. The specified random effects avoid pseudo-replication and control for variation between subjects, images and subject-specific variation of image effects with regard to the average response respectively. We included all theoretically identifiable random slopes based on at least 3 or more unique observations of the covariate per subject or at least 2 or more observations per level of the factor per subject. Thus, for ‘subject ID’, we included the interaction of valence, condition, and trial and the Image Complexity as they each meet the criteria for the specification of random slopes. For the ‘image ID’ since each image uniquely belongs to only one level of the factor valence and has one value for the image complexity rating, we could only include the interaction of valence and condition. Lastly, for the grouping factor ‘Image in subject’, each image nested within a subject belongs to only level of the factor valence and condition. Thus, we only included the random slopes of the trials for ‘Image in subject’.

The models fitted are:

GLMM -

- ***Accuracy (Valence/Old New Classification Task)*** *∼ Valence * Condition * Trials + Image Complexity + (1 + Valence*Condition* Trials + Image Complexity | subject) + (1 + Condition * Trials |Image ID) + (1 + Trials |Image in subject)*

LMM -

- ***ERP amplitudes/Response Times (Valence/Old New Classification Task)*** *∼ Valence * Condition * Trials + Image Complexity + (1 + Valence * Condition * Trials + Image Complexity |subject) + (1 + Condition * Trials |Image ID) + (1 + Trials |Image in subject)*

##### Model fitting

To keep the probability of Type 1 error at the nominal level of .05 and to avoid an overconfident model, we included all theoretically identifiable random slope components and correlations between random intercept and slope into the model. However, we excluded the correlations from the model if and when they are unidentifiable (recognizable by the majority of the absolute correlation parameters being close to 1). In addition, if we encounter convergence issues with our models, we had planned to first change the optimizers (default for LMM is ‘nloptwrap’ and for GLMM is a combination of ‘bobyqa’ and ‘Nelder_Mead’) used to fit the model. For the LMM models we fitted refit the models using ‘bobyqa’ and for the GLMM models using ‘nloptwrap’. If replacing the 23 optimizers does not help, we had planned to use the function allFit in R, which iterates through different optimizers to find the optimizer that increases the likelihood of converge. Our next planned step in dealing with convergence issues was to simplify the models, i.e., in a first step drop the random slope term for the interaction and, if convergence issues persist, drop random slope parameters whose contributions are estimated to be very small.

For the ERP data, we plotted the residuals against the fitted values and the residuals against the number of trials to assess linearity. If we find evidence of non-linearity, we had planned to initially attempt to transform the response or trial numbers to make them linear. If the transformations do not work, we had planned to then consider using a non-linear model to fit the data.

##### Model validation

For all models, the independence of the residuals and the lack of highly influential outliers was assessed. In addition, for the LMM models, we checked the assumptions of normality and homogeneity of the residuals by means of a qq-plot and residual plotted against fitted values. If any issues arise from these assumptions, we had planned to adjust the response variables accordingly by applying transformations.

We assessed the model stability using the DFBeta method by excluding levels of the grouping factor one at a time to see the effect on the model coefficients. We used the cut off ± 2/sqrt(N), where N is the sample size. If the DFBETA values are larger than the specified cutoff, we planned to report that the model estimates are unstable with respect to the specific predictor and be cautious in our interpretation of the results.

##### Transformations

The models had 2 fixed effect categorical variables that were dummy coded with “neutral valence” and “control condition” as the reference levels for the image valence and experimental conditions, respectively. Additionally, the random slopes included in the models were dummy coded and mean centered. The covariates trials and image complexity were z-transformed to make the model estimates more interpretable ^81^ and to facilitate model convergence.

The valence and old/new classification accuracies were transformed into the odds of correct or incorrect valence classification. The response time data were planned to be log-transformed to account for the original skewness of the data.

##### Inference criteria

To avoid multiple testing, we uses a full versus null model comparison using a likelihood ratio test, where the full model included both the fixed effects and the random effects, and the null model omitted all fixed effects of interest ^82^ but retained the variables and random effects to be controlled for. The fixed effects of interest are, valence, condition, and trials, and any of their interactions. To assess the significance of individual effects in the LMMs, we used the Satterthwaite approximation ^83^ using the function lmer of the package lmerTest in R and a model fitted with restricted maximum likelihood. In the case of GLMMs, assessed the contribution of each predictor using likelihood ratio tests with the drop1 method (test = Chisq).

If the full-null model comparison revealed significance, but the three-way interaction between condition, valence, trial did not, we dropped the three-way interaction, subsequently any potential non-significant two-way interactions, to obtain reliable estimates that are unconditional on particular values of other predictors with which they interact in the model.

Parametric bootstrapping was used to obtain 95% confidence intervals of the model estimates. Additionally, to interpret null findings, we used frequentist equivalence test with upper and lower bounds as the smallest meaningful effect of ±0.2.

## Results

Based on the estimated large sample size from the power analysis, we conducted an interim analysis at N = 60 and a final analysis at N = 80 to evaluate whether the data collection could be concluded earlier. To control for inflated Type 1 error due to multiple looks at the data, we applied a Pocock correction, yielding a more conservative significance threshold (alpha = 0.0294) for all the registered analyses. All LMMs were fit using the “bobyqa” optimizer, and the GLMMs used “nloptwrap”.

In our Stage 1 report, the equivalence bounds were set to ±0.2, based on the theoretical definition of small effect sizes. However, inspection of the model estimates revealed that observed effects were considerably smaller than expected. To avoid declaring practically meaningful effects as “equivalent to zero” due to overly wide bounds, the smallest meaningful effect was adjusted post hoc to ±0.1 (***equivalence tests reported in Supplementary Information 1***). This bound reflects a more conservative and context-sensitive threshold for practical relevance in this dataset, aligning better with the empirical distribution of effects. Further, the equivalence boundary aligns with the confidence intervals reported for P1 amplitudes in prior associative learning studies ^84^.

Since the registered models included three random-effects factors and their corresponding random slopes, we removed the correlations between slopes and intercepts across all models to reduce model complexity and improve model efficiency.

We had also preregistered the use of DFBeta diagnostics to assess model stability by iteratively removing each level of the grouping factor (i.e., participants) and examining the impact on model coefficients. However, this approach proved infeasible due to the computational burden and extended runtimes involved in refitting the full mixed-effects models for each of the 80 participants. Our models included multiple random effects and interaction terms, and convergence issues were frequently encountered during even a single model fit, making iterative refitting impractical and unreliable. As a practical alternative, model stability was evaluated in the explorative analyses using data averaged across trials (***model stability values are reported in the results table in Supplementary Information 2***).

### Valence Classification Task

To assess whether our experimental setup successfully induced perceptual uncertainty, we examined mean accuracy across the first two blocks for each participant in the social and control conditions. The majority of participants (N = 63) exhibited higher accuracy in the social condition compared to the control condition, indicating their reliance on social cues to resolve perceptual uncertainty. However, a subset of participants (N = 17) performed better in the control condition than in the social condition, potentially reflecting individual differences in strategy use or a lack of trust in the social cue within a lab-based artificial social learning paradigm. These participants may have perceived the facial expressions as unreliable or irrelevant, opting instead for their own subjective valence judgments.

Additionally, participants often defaulted to a ‘neutral’ response when highly uncertain, which may have masked condition differences at the individual level. This response bias is reflected in the relatively high frequency of neutral classifications for both positive and negative target images – even in the social condition, where participants frequently chose the neutral option despite the presence of facial expressions (social: N = 6059; control: N = 9300). Despite individual variability in the valence classification task, the planned statistical model revealed significantly higher accuracy for emotional target images in the social compared to the control condition. This suggests that most participants incorporated the social cues into their decision-making. Based on this overall pattern, no participants were excluded from subsequent analyses based on their initial performance in the first two blocks.

#### Valence Classification Accuracy and RT (Fig. 2a & 2b)

**Figure 2.**
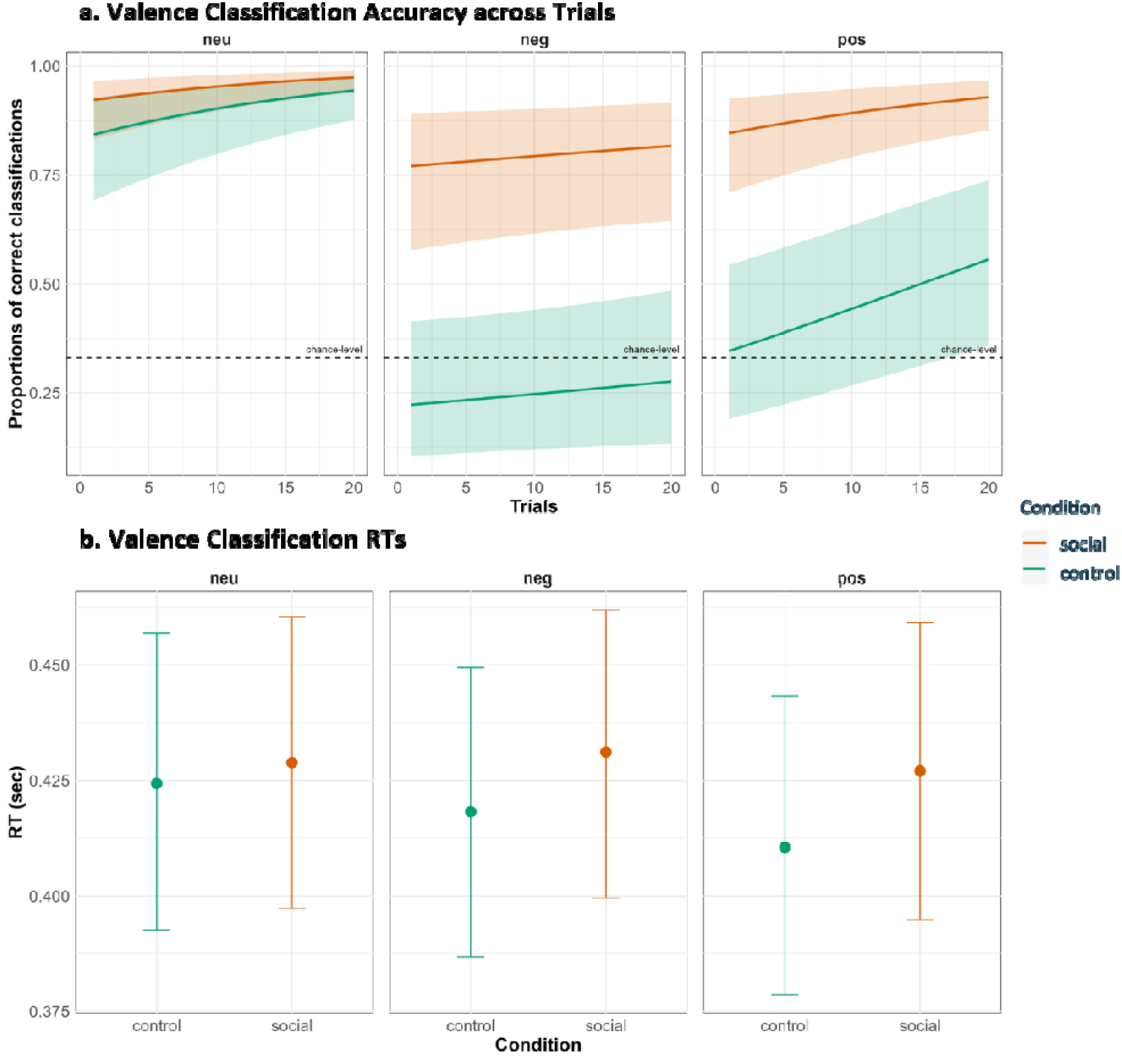
a. Valence Classification Accuracy across Trials. b. Valence Classification RTs aggregated across Trials. Plots indicate mean model-based predicted values along with the corresponding 95% CIs.

A significant valence × condition interaction emerged for classification accuracy. Relative to neutral target images, accuracy was significantly higher in the social compared to the control condition for both negative and positive target images (**negative:** β = 1.667, SE = 0.425, z = 3.919, p < .001; **positive:** β = 1.551, SE = 0.421, z = 3.684, p < .001). For response times (RTs), we also observed a valence × condition interaction, again relative to neutral target images; however, the effect was contrary to expectations. Rather than faster responses, the presence of a social cue was associated with marginally (p-value made conservative after Type 1 correction) longer RTs for positive target images (**positive:** β = 0.012, SE = 0.006, t = 2.111, p = .040).

While overall performance improved across trials, the model revealed no significant condition × trial interactions for either valence classification accuracy or RTs, contrary to our predictions. However, a significant valence × trial interaction emerged for accuracy for the negative target images, where accuracy gains over time were reduced compared to neutral ones (**negative:** β = -0.262, SE = 0.088, z = - 2.972, p = .003). This pattern suggests a reduced perceptual learning effect for negative target images. For RTs, a main effect of trial was observed, with RTs decreasing across trials regardless of valence or condition (**trial:** β = −0.045, SE = 0.005, t = −9.593, p < .001), indicating general task-related learning and improved processing efficiency over time.

To assess whether participants formed associations between the target images and their respective conditions, we administered recall and recognition memory tasks immediately after the learning session. Performance on both tasks exceeded chance levels (recall: 25%; recognition: 50%), indicating successful learning across participants. In the recall task (**Fig. S1a – see Supplementary Information 1**), mean accuracy was higher for target images in the social condition (67.98%) than in the control condition (59.27%), suggesting enhanced memory encoding when social cues were present. In contrast, recognition accuracy (**Fig. S1b - see Supplementary Information 1**) was comparable between the social (72.20%) and control (72.13%) conditions, indicating no condition-specific differences in recognition memory.

### ERP amplitudes – Valence Classification Task (Learning Session)

#### P1 (Fig. 3a)

**Figure 3.**
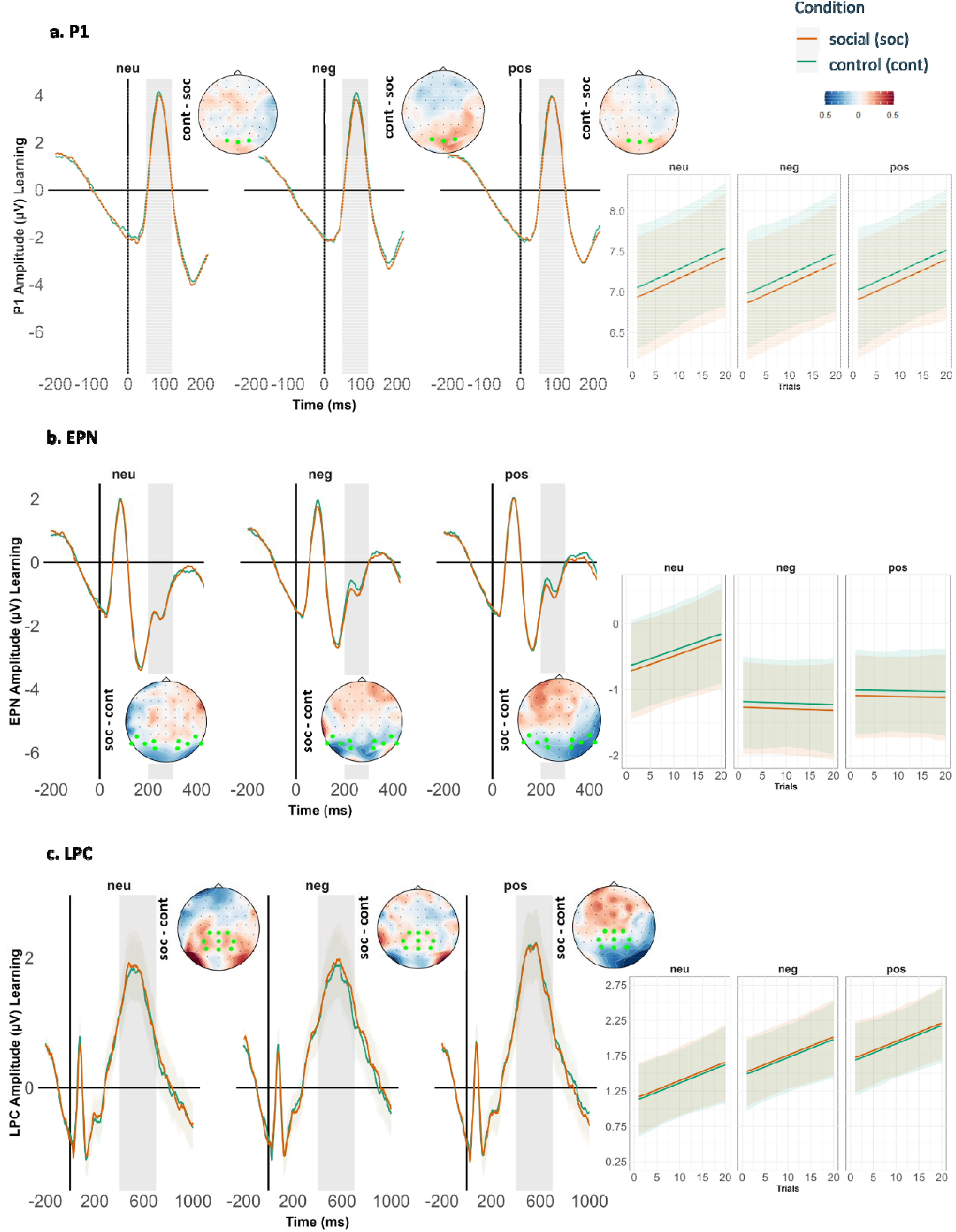
a. P1. b. EPN. c. LPC. Grand average waveforms across conditions for each of the target image valence categories (neu, neg, pos). Gray rectangular shaded region marks the time-window of the ERP. Topoplots indicate difference between conditions and green dots mark the corresponding ROIs for each of the ERPs. The corresponding line plots represent mean model-based predicted value across trials with shaded regions indicating 95% CIs.

Due to substantial inter-individual variability in P1 onset (ranging from around 50 to 120 ms), we identified the peak amplitude within this window for each participant and trial. To obtain a stable estimate, mean amplitude was calculated by averaging across a ±10 ms window around the individual peak. These single-trial P1 amplitudes from the relevant ROIs were entered into a LMM.

The comparison between the full model and a null model (including only control variables and random effects) was not statistically significant, suggesting that the inclusion of fixed effects did not improve model fit (χ^2^ = 8.880, df = 4, *p* = .064). Contrary to our hypothesis, the main effect of condition – although not significant – was in the opposite direction: P1 amplitudes were numerically lower in the social condition compared to the control condition (**condition:** β = −0.118, SE = 0.065, *t* = −1.804, *p* = .079). No significant interactions involving condition, valence, or trial were observed. However, a significant main effect of trial emerged, with P1 amplitudes increasing over time, irrespective of condition or valence (**trial:** β = 0.136, SE = 0.057, *t* = 2.389, *p* = .020).

#### EPN (Fig. 3b)

Single-trial EPN amplitudes were extracted from the relevant ROIs within the 200–300 ms time window and analyzed using a LMM. Contrary to predictions, the model revealed no significant effect of condition on EPN amplitudes (**condition:** β = −0.086, SE = 0.061, *t* = −1.409, *p* = .166), indicating that the presence of a social cue did not modulate early emotional attention during the learning session. Instead, EPN amplitudes were significantly modulated by a valence × trial interaction: across trials, both negative and positive (**negative:** β = −0.146, SE = 0.056, *t* = −2.585, *p* = .012; **positive:** β = −0.141, SE = 0.062, *t* = −2.255, *p* = .028) target images elicited stronger EPN responses relative to the neutral target images.

#### LPC (Fig. 3c)

Single-trial LPC amplitudes were extracted from the relevant ROIs within the 400–600 ms time window and analyzed using a LMM. Consistent with the EPN results, the model revealed no significant effect of condition (**condition:** β = 0.035, SE = 0.050, *t* = 0.697, *p* = .490), suggesting that the presence of a social cue did not modulate later-stage evaluative processing during learning. Contrarily, LPC amplitudes were significantly modulated by main effects of valence and trial. Specifically, amplitudes were overall enhanced for positive target images (**positive:** β = 0.555, SE = 0.191, *t* = 2.905, *p* = .005) and increased across trials, irrespective of valence or condition (**trial:** β = 0.136, SE = 0.053, *t* = 2.588, *p* = .012).

### Old/New Classification Task

#### Old/New Classification Accuracy and RT (Fig. S2a & S2b – see Supplementary Information 1)

Old/New classification accuracy was significantly modulated by a three-way interaction between valence, condition, and trials. For positive target images, accuracy increased more steeply across trials in the social condition compared to the control condition, relative to the difference between conditions for neutral target images (**positive:** β = 0.745, SE = 0.301, *z* = 2.479, *p* = .013). However, interpretation of this three-way interaction is limited by ceiling effects, as recognition accuracy for positive and neutral target images was already high across conditions, reducing sensitivity to subtle condition differences. Thus, the observed interaction may reflect compressed variance rather than genuine improvements in memory retrieval over time. Old/New Classification RTs were significantly influenced by trial, with faster classification responses across repeated exposures (**trial:** β = −0.048, SE = 0.004, *t* = -12.750, p < .001). As predicted, no effect of condition on RTs was observed (**condition:** β = 0.000, SE = 0.001, *t* = -0.003, p = .999).

### ERP amplitudes – Old/New Classification Task (Test Session)

#### P1 (Fig. 4a)

**Figure 4.**
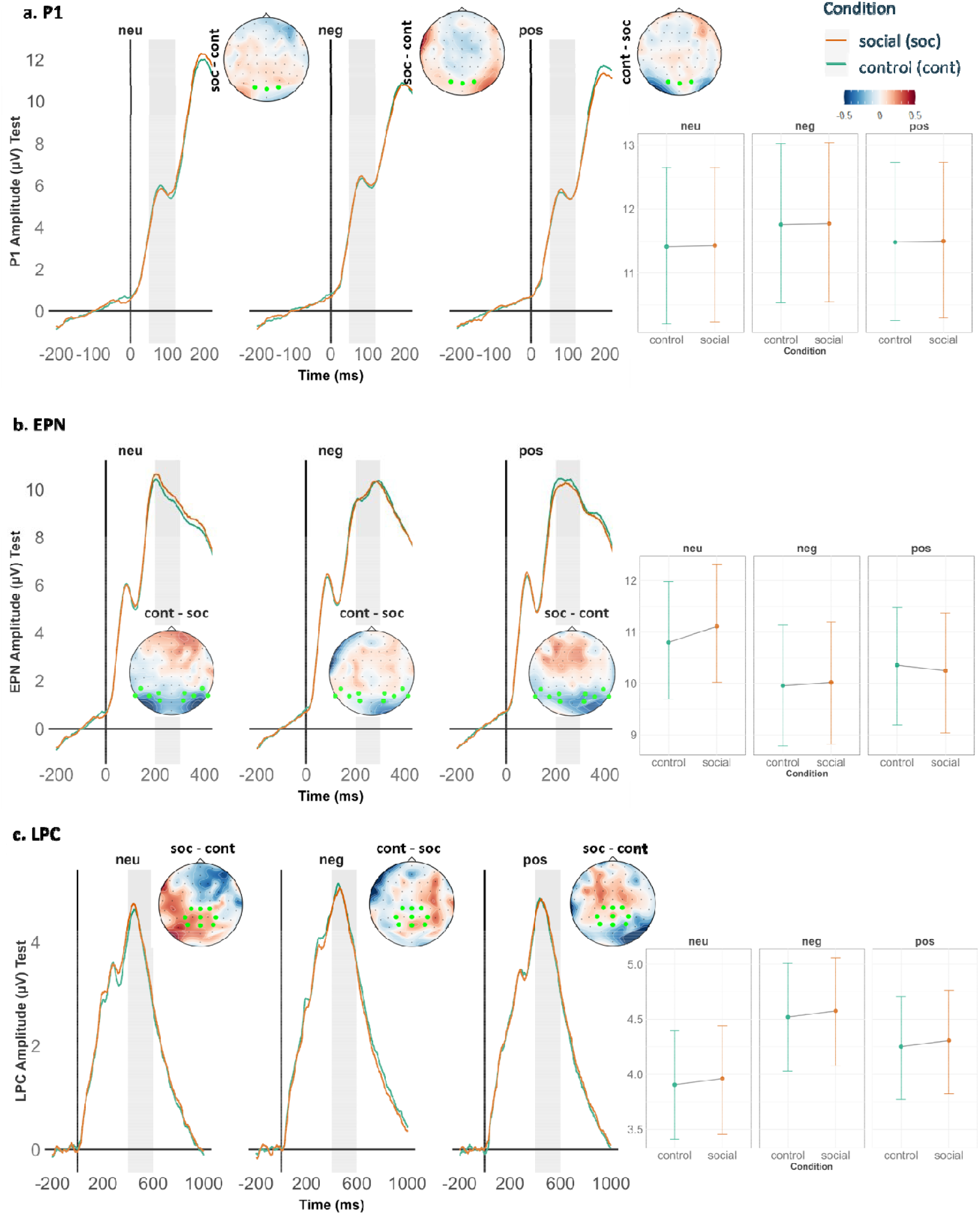
a. P1. b. EPN. c. LPC. Grand average waveforms across conditions for each of the target image valence category (neu, neg, pos). Gray rectangular shaded region marks the time-window of the ERP. Topoplots indicate difference between conditions and green dots are the corresponding regions of interest (ROIs) for each of the ERPs. The corresponding line plots represent mean model predicted values aggregated across trials with error bars regions indicating 95% CIs.

Due to substantial variability in P1 onset across participants (ranging from 50 to 120 ms), we identified the peak amplitude within this window for each participant and trial. To obtain a stable estimate, mean amplitudes were calculated by averaging over a ±10 ms window around the peak. These single-trial P1 amplitudes, extracted from the relevant ROIs, were entered into a LMM.

P1 amplitudes were significantly modulated by a main effect of trial, with amplitudes increasing over time (**trial:** β = 0.365, SE = 0.071, t = 5.152, p < .001). In contrast to our predictions, no significant effect of condition was observed (**condition:** β = 0.015, SE = 0.073, t = 0.211, p = .746).

#### EPN (Fig. 4b)

Single-trial EPN amplitudes, extracted from the relevant ROIs within the 200–300 ms time window, were analyzed using a LMM. EPN amplitudes were significantly modulated by a valence x condition interaction, with more pronounced amplitudes for positive target images in the social condition compared to the control condition, relative to the difference between conditions for neutral target images (**positive:** β = -0.422, SE = 0.172, t = -2.453, p = .017).

Additionally, a marginal (p-value made conservative after Type 1 correction) valence x trial interaction emerged across both conditions, indicating that emotional target images consistently elicited stronger EPN amplitudes across trials relative to neutral target images (**negative:** β = -0.148, SE = 0.072, t = -2.070, p = .044; **positive:** β = -0.151, SE = 0.069, t = -2.195, p = .033).

#### LPC (Fig. 4c)

Single-trial LPC amplitudes, extracted from the relevant ROIs within the 400–600 ms time window, were analyzed using a LMM. LPC amplitudes were modulated by an interaction of valence x trials, with marginally (p-value made conservative after Type 1 correction) significant extinction effects observed for negative target images across trials, relative to the neutral target images (**negative:** β = - 0.132, SE = 0.062, t = -2.111, p = .040). No significant effect of condition emerged (**condition:** β = 0.057, SE = 0.050, t = 1.126, p = .263).

#### Old/New Effects

To examine the Old/New effect—typically characterized by more positive ERP amplitudes for previously seen (“old”) images compared to unseen (“new”) images in the P300 and LPC time windows—we conducted LMM analyses. Due to missing triggers for some new images, trial-wise EEG data could not be extracted. Thus, the models were fitted using average amplitudes for each valence-by-condition combination. Analyses were restricted to old images in the control condition, which were then compared to new images.

#### P300 (Fig. S3a – see Supplementary Information 1)

As expected, old images elicited significantly greater positive amplitudes than new images (old: β = 0.861, SE = 0.071, t = 12.065, p < .001). This old/new effect was more pronounced for emotional target images relative to neutral images (negative: β = 1.107, SE = 0.087, t = 12.667, p < .001; positive: β = 0.244, SE = 0.071, t = 3.425, p = .005).

#### LPC (Fig. S3b – see Supplementary Information 1)

Similarly, LPC amplitudes were significantly larger for old than new images (**old:** β = 0.660, SE = 0.057, t = 11.501, p < .001), with greater enhancement for emotional target images compared to neutral images (**negative:** β = 0.709, SE = 0.070, t = 10.082, p < .001; **positive:** β = 0.287, SE = 0.070, t = 4.084, p < .001).

## Exploratory Results

Because target images were presented under conditions of perceptual uncertainty, participants often relied on their own subjective valence judgments of the target images. Consequently, some target images were classified in accordance with their normative valence (i.e., accurately), while others were misclassified (i.e., inaccurately). When subjective valence judgments were taken into account, both behavioral and ERP measures reflected whether participants’ subjective evaluations aligned or diverged from the normative valence of the target images. For the learning session, we excluded 6.3 % of trials (3480 data points) in which participants made a directly opposing classification – for example, labeling a negative target image as positive or vice versa. Accordingly, for normatively positive or negative target images, “inaccurate classifications” included only trials where participants responded with a “neutral” classification. For normatively neutral target images, inaccurate classifications encompassed both “positive” and “negative” responses. For the learning session, every trial was coded separately as ‘accurate’ or ‘inaccurate’ classification based on the match between the subjective valence judgment and the normative image valence. For the test session, since the task did not necessitate valence judgments, the classification accuracies were based on the learning session performance. Accordingly, the target images in the test session were assigned a classification accuracy score reflecting the frequency of consistent classifications during the learning session. Specifically, for each target image, we identified the most frequently selected classification valence over the final 18 of its 20 presentations (the last 18 presentations were used to reduce noise as participants may have been more indecisive during the initial two presentations while adjusting to the task). If this most frequent valence matched the normative value on at least 14 occasions, the image was classified as an ‘accurate’ classification; otherwise, it was classified as an ‘inaccurate ‘classification. All the exploratory analyses were conducted using data averaged across trials per participant and condition, for both behavioral and ERP measures. For the factor ‘classification accuracy’, “accurate” was used as the reference level.

While perceptual uncertainty was experimentally induced, including classification accuracy in the models allowed us to account for subjective uncertainty, as the experimental design introduced considerable inter-individual differences in classification strategies.

### Valence Classification Task – Learning Session

#### RTs accounting for classification accuracy (Fig. 5)

**Figure 5.**
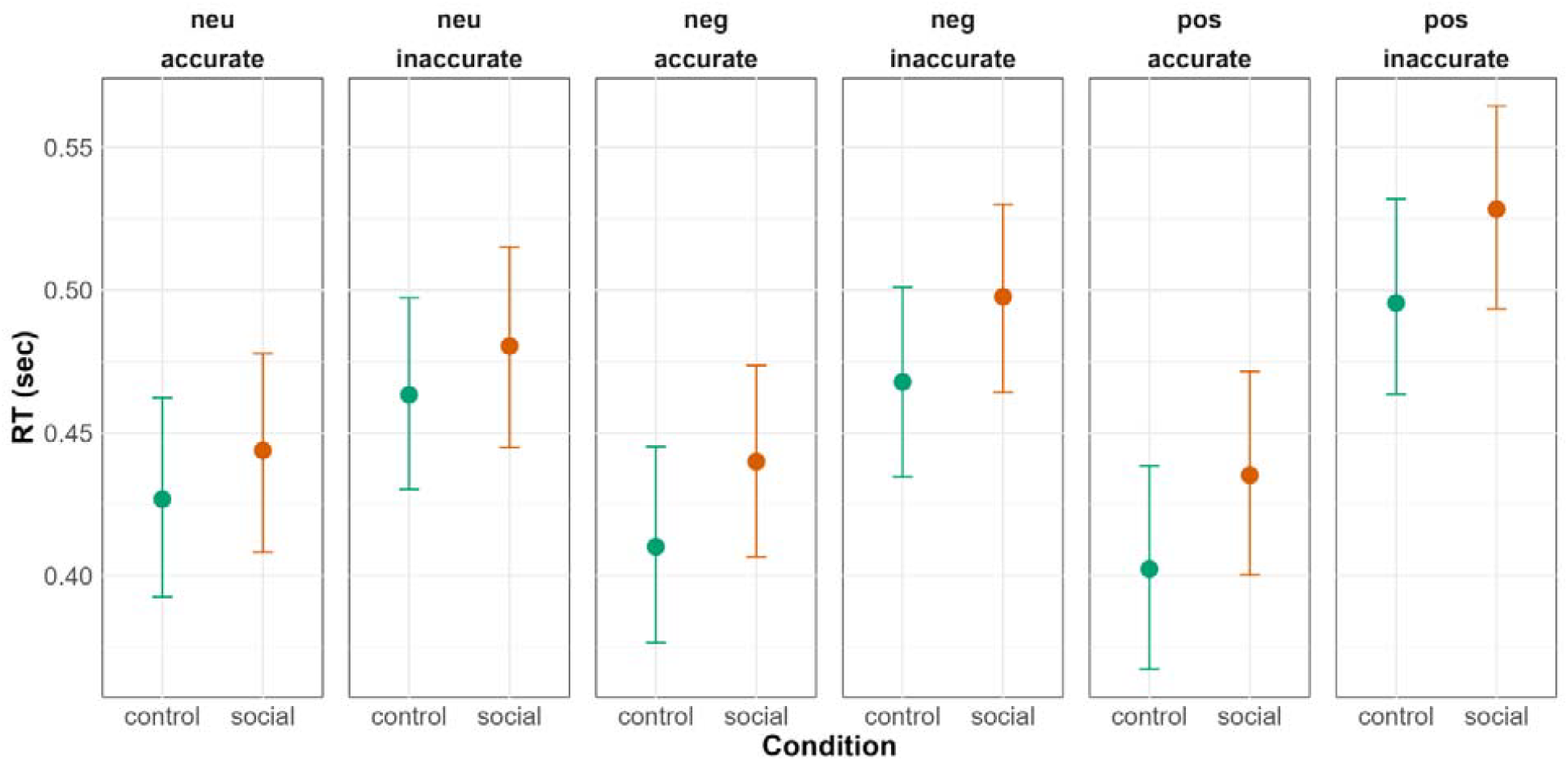
Valence Classification RTs split by classification accuracy. Plots represent mean model-based predicted values with their corresponding 95% CIs as error bars.

RT analyses revealed a valence × classification accuracy interaction. Specifically, inaccurately classified positive target images – those judged as neutral—elicited significantly longer RTs relative to the difference between conditions observed for the neutral target images (**positive:** β = 0.057, SE = 0.014, *t* = 4.041, *p* < .001). A similar trend was observed for inaccurately classified negative images, but this effect did not reach statistical significance (**negative:** β = 0.021, SE = 0.015, *t* = 1.416, *p* = .163).

#### P1 Amplitudes Accounting for Classification Accuracy (Fig. 6a)

**Figure 6.**
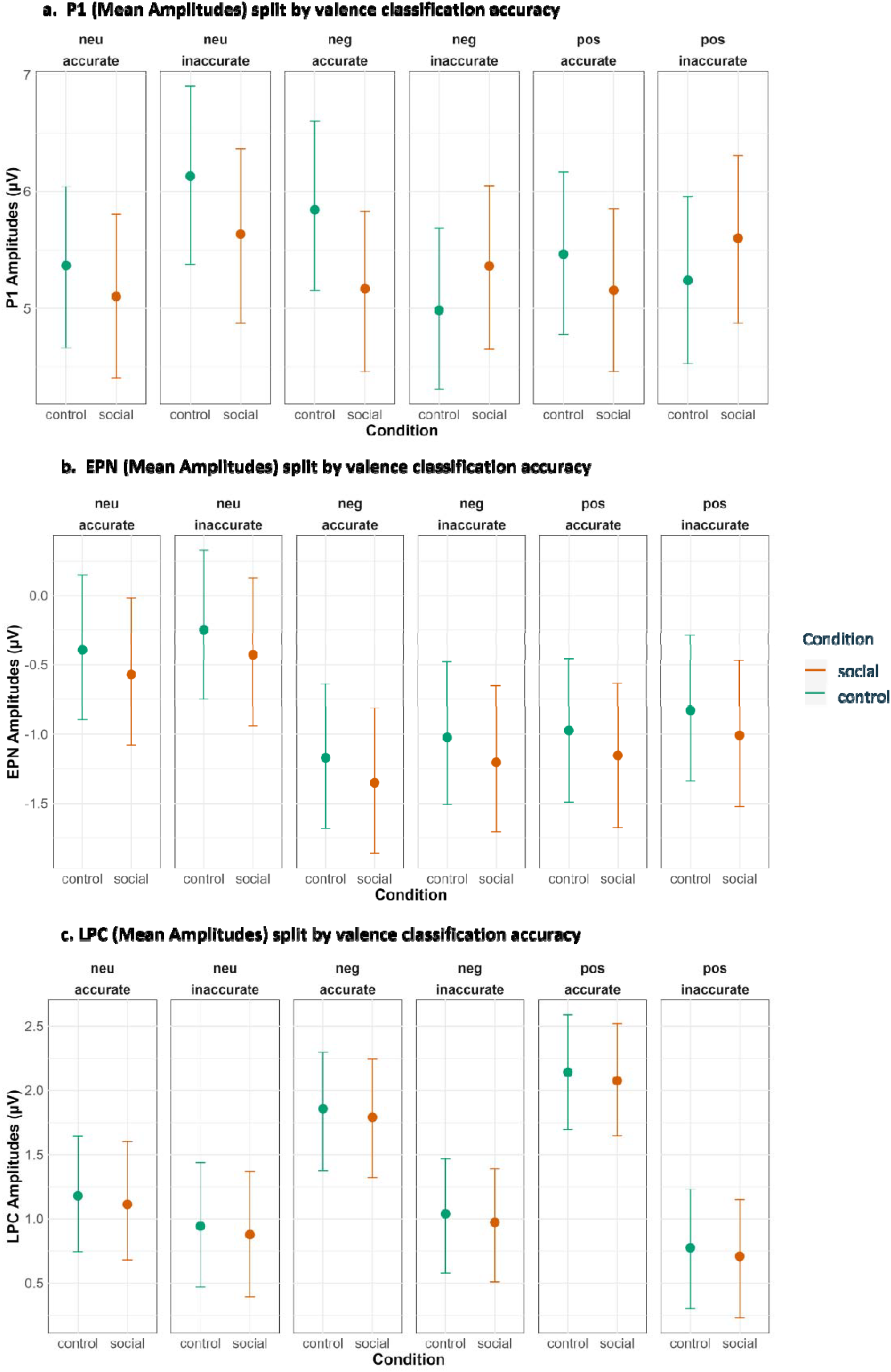
Mean ERP Amplitudes from corresponding windows and regions of interest (ROIs) a.P1 b. EPN c. LPC. ERPs split by classification accuracy. Plots represent mean model-based predicted values with their corresponding 95% CIs as error bars.

P1 amplitudes, referenced to neutral target images, were significantly modulated by a three-way interaction between valence, condition, and classification accuracy for negative target images (**negative:** β = 1.280, SE = 0.468, *t* = 2.737, *p* = .006), and showed an effect in the similar direction – however not significant - for positive target images (**positive:** β = 0.901, SE = 0.479, *t* = 1.877, *p* = .060). For neutral target images, P1 amplitudes were lower in the social compared to the control condition, regardless of classification accuracy. For negative target images, P1 amplitudes were similarly reduced in the social condition compared to the control when classifications were accurate. However, this pattern reversed for inaccurately classified negative images, where P1 amplitudes were enhanced in the social condition relative to the control condition.

#### EPN Amplitudes Accounting for Classification Accuracy (Fig. 6b)

EPN amplitudes were significantly modulated by normative valence, with enhanced amplitudes for both negative and positive target images compared to neutral images. (**negative:** β = -0.778, SE = 0.146, t = -5.318, p < .001; **positive:** β = - 0.582, SE = 0.145, t = -4.023, p < .001). No significant effects of classification accuracy were observed. However, when accounting for classification accuracy, a trend emerged toward greater EPN amplitudes in the social compared to the control condition (β = -0.180, SE = 0.098, t = -1.840, p = .069).

#### LPC Amplitudes Accounting for Classification Accuracy (Fig. 6c)

LPC amplitudes were significantly modulated by a valence x classification accuracy interaction: Specifically, inaccurately classified emotional target images elicited reduced amplitudes compared to accurately classified ones, relative to the pattern observed for neutral target images (**negative :** β = -0.582, SE = 0.205, t = -2.838, p = .006 ; **positive :** β = -1.131, SE = 0.226, t = -4.996, p < .001).

### Old/New Classification Task –Test Session

#### Old/New Classification Accuracy Accounting for Classification Accuracy (Fig. S4 – see supplementary Information 1)

Old/New classification accuracy was modulated by an interaction between valence x classification accuracy, relative to neutral images. Specifically, emotional images that had been inaccurately classified on Day 1 were significantly less likely to be remembered than accurately classified images (**negative:** β = -1.503, SE = 0.240, z = -6.276, p < .001; **positive:** β = -1.461, SE = 0.255, z = -5.734, p < .001).

#### EPN Amplitudes Accounting for Classification Accuracy (Fig 7a)

**Figure 7.**
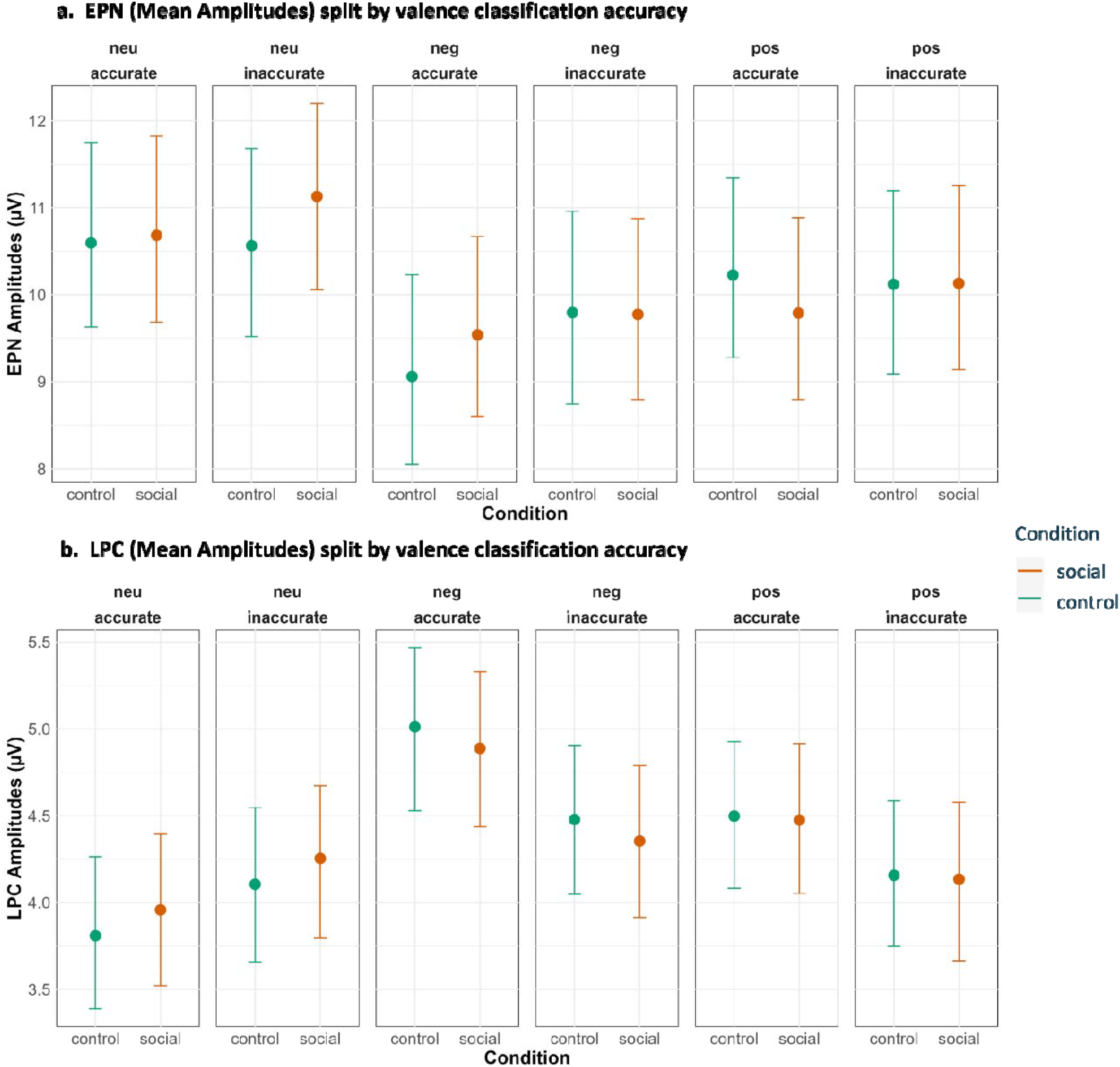
Mean ERP Amplitudes from corresponding windows and regions of interest (ROIs) a. EPN b.LPC. ERPs split by classification accuracy. Plots represent mean model-based predicted values with their corresponding 95% CIs as error bars.

EPN amplitudes, in reference to the neutral target images, were significantly modulated by a three-way interaction between valence, condition, and classification accuracy for the negative target images (**negative :** β = -0.978, SE = 0.470, t = -2.082, p = .040). To disentangle these effects, separate models were fitted for accurately and inaccurately classified images.

For inaccurate classifications, EPN amplitudes for neutral target images were less pronounced in the social condition compared to the control condition, In contrast, EPN amplitudes for both negative and positive target images were enhanced but comparable to those in the control condition (**negative:** β = - 0.692, SE = 0.340, t = -2.034, p = .045; **positive:** β = -0.753, SE = 0.359, t = -2.098, p = .039). For accurate classifications, EPN amplitudes for neutral images remained comparable across conditions. However, for negative target images, the social condition elicited numerically – but not significantly – smaller EPN amplitudes compared to the control condition (**negative:** β = 0.282, SE = 0.319, t = 0.882, p = .380). In contrast, accurately classified positive target images continued to elicit enhanced EPN amplitudes in the social compared to the control condition (**positive:** β = -0.638, SE = 0.268, t = -2.382, p = .018).

#### LPC Amplitudes Accounting for Classification Accuracy (Fig. 7b)

LPC amplitudes were significantly modulated by a valence x condition interaction for negative target images. Specifically, while amplitudes were comparable between conditions for neutral target images, they were reduced in the social compared to the control for negative target images (**negative:** β = -0.272, SE = 0.132, t = -2.056, p = .043). Consistent findings from the learning session, LPC amplitudes were also influenced by a valence x classification accuracy interaction. Relative to the pattern observed for neutral target images, inaccurate classifications of emotional target images led to reduced LPC amplitudes compared to accurate classifications (**negative:** β = -0.830, SE = 0.145, t = -5.723, p < .001; **positive:** β = -0.637, SE = 0.144, t = - 4.429, p < .001).

## Discussion

The present study examined whether emotional salience can be transferred from social cues – specifically, facial expressions – to ambiguous visual stimuli under conditions of experimentally induced perceptual uncertainty. During the learning session, participants performed a valence classification task in which some images were paired with emotional facial expressions (social condition) while others were paired with scrambled versions of faces that carried no emotional content (control condition). On the following day, preservation of emotional salience in memory was tested in an Old/New classification task.

We registered the hypothesis that emotional salience would be transferred from facial expressions to target images during learning. Specifically, we expected condition-specific modulations of early (P1), mid (EPN), and late (LPC) ERP components over repeated exposures to reflect this associative process. However, these predictions were not supported. ERP amplitudes during the learning session did not differ by condition, and equivalence tests confirmed that the observed effects were statistically equivalent to zero within predefined bounds. To interpret these null findings, we introduce a post hoc theoretical framework that emphasizes the role of subjective uncertainty in modulating social cue influence. This metacognitive perspective, though not preregistered, offers a plausible explanation for when and how social information is integrated into learning.

Relative to the neutral target images, social cues were significantly used for valence classification of emotional target images. However, ERP evidence did not indicate emotional salience transfer across trials as predicted. Instead, the influence of social cues during the learning session appeared to vary depending on participants’ metacognitive state of subjective uncertainty as reflected in the P1 modulations in response to negative target images. Recognition memory after overnight consolidation showed evidence of additive emotional salience from the social cues at the EPN level for accurately classified positive target images in comparison to the neutral target images. In later stages of processing, LPC amplitudes were reduced for negative target images in the social condition, irrespective of classification accuracy. Together, our findings suggest that the influence of social cues is contingent on the metacognitive state of subjective uncertainty and does not automatically induce emotional salience transfer during learning. This pattern became emergent in the test session after overnight consolidation: Positive target images – associated with lower subjective uncertainty – showed enhanced emotional salience, while negative target images, marked by higher uncertainty, led to greater reliance on social cues and weaker encoding of the image itself.

### Day 1: Learning Session - Valence Classification Task

As predicted, social cues significantly improved valence classification accuracy, particularly for emotional target images. This finding aligns with previous findings suggesting that social information becomes especially valuable under conditions of uncertainty, ambiguity, or when individuals seek external guidance for decision-making ^59,4,60,67,61^. However, contrary to our expectations, these accuracy benefits did not increase over trials. Instead, both classification accuracy and RTs improved similarly across social and control conditions – suggesting a general effect of perceptual learning. Previous studies have demonstrated trial-wise calibration of advice or social inputs leading to cumulative learning ^30,85^. In contrast, our paradigm offered limited incentives for such calibration: target images were repeatedly paired with the same social cue, and participants did not receive feedback about their classification accuracy. Moreover, the inclusion of a ‘neutral’ response option allowed participants to indicate subjective uncertainty explicitly – even when a social cue was available—potentially diluting any consistent reliance on external cues over time.

A general trend of slower RTs in the social condition compared to the control condition for positive target images support the interpretation that social cues were used selectively based on participants’ metacognitive state. This result deviates from our preregistered prediction that social cues would reduce uncertainty and thereby facilitate faster responses. Instead, the prolonged RTs likely reflect additional cognitive demands, with participants engaging in more deliberate evaluative processes – potentially weighing their subjective uncertainty against the valence information provided by the social cue. Consistent with this interpretation, exploratory analyses accounting for classification accuracy as a proxy for subjective uncertainty showed slower responses for inaccurate compared to accurate classifications, particularly for positive target images, suggesting that accurate classifications were associated with greater certainty than inaccurate ones.

Neural modulation during early perceptual processing, as indexed by the P1 ERP component, did not support the predicted transfer of emotional salience from social cues to the ambiguous target images. Previous studies indicated that social cues can modulate early attentional processes, particularly under conditions of uncertainty or social pressure ^69,70,66,71,68^. In line with this, our data revealed metacognitive state-dependent modulations of the P1 component.

The registered models for P1 indicated a pattern for reduced amplitudes in the social compared to the control condition, however accounting for classification accuracy as a proxy for subjective uncertainty helped tease apart the P1 modulations better. For neutral target images, P1 amplitudes were reduced across both social and control conditions when classifications were accurate, consistent with low subjective uncertainty and reduced perceptual demands for neutral target images. When participants misclassified neutral images as emotional, P1 responses diverged: amplitudes increased in the control condition—indicating heightened sensory engagement under uncertainty—but decreased in the social condition, suggesting that participants used the facial expression to resolve uncertainty and suppress early visual processing.

A complementary pattern emerged for emotional target images, particularly negative ones. Behavioral findings showed weaker perceptual learning across trials for negative compared to neutral target images, suggesting less stable internal judgments and a greater reliance on social cues. This was mirrored in P1 amplitudes: when negative images were accurately classified in the social condition, amplitudes were reduced compared to control, suggesting facilitated classification through social cue alignment. Conversely for inaccurately classified emotional images, P1 amplitudes were increased in the social condition, indicating greater sensory engagement under uncertainty – internal and external signals conflicted.

These findings are consistent with predictive coding models ^86,87^, which propose that prior information – including social cues – modulates early sensory processing based on the magnitude of the prediction error. When internal expectations and external cues align, prediction error is low and sensory responses are suppressed; when they conflict, prediction error is high, prompting greater sensory engagement. Thus, rather than directly shaping early perceptual representations, social cues appear to exert a flexible, metacognitive influence – either dampening or enhancing early visual responses depending on subjective uncertainty ^4,60,67,61^.

No evidence was found for the predicted transfer of emotional salience from social cues to target images at mid or late stages of processing, as indexed by the EPN and LPC components. Instead, both ERP markers were modulated by emotional valence regardless of cue condition.

The EPN amplitudes in both the registered and the exploratory models showed significantly greater amplitudes for emotional compared to neutral target images, consistent with its established role in bottom-up attentional prioritization of emotionally salient stimuli across different stimulus domains ^46–51^. The absence of any condition effect suggests that emotional salience may have been driven primarily by intrinsic stimulus properties, with limited added value from the social cues. This aligns with previous findings that EPN modulations are by low-level perceptual features and are often insensitive to context or task demands ^88,89,48^. Although we controlled for image complexity in our analyses, it remains possible that residual perceptual differences contributed to the observed valence effects—particularly relevant given the experimentally induced perceptual uncertainty in our task.

The LPC component revealed robust effects of classification accuracy: amplitudes were consistently higher for correctly classified emotional images than for misclassified ones. This finding supports the interpretation of the LPC as a marker of sustained evaluative processing and affective memory encoding ^53,53,52,46,48,54,49,55^. Notably, LPC amplitudes were unaffected by social cue condition, suggesting that late-stage emotional evaluation was driven primarily by internal decision certainty rather than external cue influence.

Previous studies have shown that inherently neutral stimuli—such as pseudowords or neutral faces—can acquire motivational salience through associations with gain or losses, as reflected in EPN and LPC modulations ^40–43,90,91^. In those paradigms, participants lacked prior affective evaluations of the stimuli, which likely facilitated stronger and more consistent associative effects. In contrast, the target images in our study allowed for subjective valence judgments, which may have influenced how social cues were interpreted and used. Rather than forming new associations, participants may have used facial expressions as secondary inputs in the appraisal process—adjusting or affirming their initial judgments depending on their level of subjective uncertainty. Future research could test this possibility more directly by restricting stimulus content to truly neutral items and assessing whether emotional social cues can more reliably induce associative salience under such conditions.

### Day 2: Test Session – Old/New Classification Task

Recognition accuracy for emotional target images was significantly modulated by the classification accuracy during the learning session. Specifically, target images that were correctly classified during the learning session on Day 1 were more likely to be recognized in the test session on Day 2. This suggests that successful encoding of emotional content – whether based on internal judgments or supported by cues –facilitated subsequent memory, consistent with well-established links between affective salience and memory ^92–95^. Importantly, no additional memory benefit emerged for target images paired with social cues. This aligns with the ERP results from the learning session, where LPC amplitudes were driven by classification accuracy rather than cue condition, indicating that memory formation was more depended on the strength of subjective emotional evaluation than on the presence of social cues.

In contrast to the P1 modulations observed during the learning session, no significant differences in P1 amplitudes were found during the test session – neither as a function of condition nor valence. Although the P1 component is often associated with early visual attention, its sensitivity to emotional content has been inconsistent across studies. While some have reported enhanced P1 responses for emotional stimuli (e.g., ^96,97^, others have failed to find reliable effects of emotion on the P1 (e.g. ^58,90,98^. Our findings align with the latter, suggesting that neither emotional valence nor prior social association influenced early perceptual processing.

This result diverges from previous associative learning studies, where P1 modulations were observed for stimuli paired with motivational outcomes – even when those incentives were no longer present during subsequent old/new recognition tasks ^40–42,99,43^. However, other studies have also failed to observe early ERP effects of learned salience under similar test conditions, particularly when associations were formed in cross-modal paradigms or involved socially or affectively complex cues ^100,98^.

In contrast to the absence of condition effects on the EPN during the learning session, the registered model for the Day 2 test session revealed enhanced EPN amplitudes for the positive images in the social condition, relative to the control. Accounting for classification accurately, the effect was primarily driven by the accurately classified positive target images. No such effect was found for negative target images. This condition-specific effect, measured relative to the difference observed between conditions for neutral images, suggests that positive images previously paired with social cues exerted a lasting influence on early emotional attention, even when the social cue was no longer present and the task did not require emotional classification.

These findings are consistent with Schacht & Vrtička, 2018 ^101^ who showed that positive emotional images depicting social content elicited enhanced EPN amplitudes, effectively counteracting the typical mid-stage negativity bias. In their study, a significant interaction between emotional valence and social content of the stimuli revealed that positive social stimuli elicited stronger attentional engagement, whereas no such enhancement was found for negative images ^102^.

Further support for a temporal dissociation between positive and negative motivational outcomes comes from Grassi et al., 2024 ^103^, who found that loss-associated stimuli influenced ERP responses during the initial learning phase, while gain-associated stimuli modulated EPN amplitudes only during a later test phase. These findings suggest that losses primarily shape early encoding processes, while gains exert stronger effects at retrieval ^42^. A similar temporal dynamic is evident in our results: negative social cues appeared to influence early perceptual classification (reflected in P1 modulations), whereas only positive social cues enhanced emotional processing after memory consolidation, as indexed by elevated EPN amplitudes during Day 2. This temporal dissociation highlights the distinct neural trajectories by which positive and negative social information contribute to affective learning and memory.

Positive social cues may have been particularly effective because of their inherently rewarding nature. Prior studies suggest that happy facial expressions function as social reinforcers, akin to primary rewards like money ^104,90,105^, and are often interpreted as broadly rewarding rather than as specific valence signals. This interpretation is supported by activation of reward-related brain areas, such as the nucleus accumbens and the medial prefrontal cortex, in response to positive stimuli ^106^. The observed EPN enhancement for accurately classified positive images in the social condition may reflect a broadened attentional state during encoding – consistent with the broaden-and-build theory of positive emotion ^107,108^. By expanding attentional scope and supporting richer associative processing, positive social cues likely enhanced consolidation and promoted prioritized reactivation during later retrieval ^109^.

In contrast to the additive salience effects for positive target images at the EPN level, negative target images elicited reduced LPC amplitudes in the social compared to the control condition. This suggests a different encoding dynamic: participants may have relied more heavily on social cues when classifying negative images, likely due to weaker or less accessible internal valence representations. Early P1 modulations support this interpretation, pointing to social cue-driven perceptual facilitation. However, such reliance on external cues may have come at the cost of deeper stimulus processing. From a metacognitive perspective, heightened uncertainty may have shifted attention away from the image itself, resulting in attenuated elaborative encoding and weaker memory traces—consistent with prior work showing that negative affect can amplify early processing while dampening later task-related stages ^110^. In contrast, for positive target images, stronger internal signals that aligned with the social cue likely supported deeper encoding and greater emotional salience at retrieval.

While previous research on social appraisal has demonstrated that emotional evaluations of stimuli can be shaped by concurrent social cues—often through expectation-driven or perspective-taking mechanisms ^33,34,20,21^ these effects have typically been studied in single-trial designs and in the presence of a social agent. In contrast, our study examined whether emotional salience could be transferred to stimuli through repeated co-occurrence with emotional expressions and persist in the absence of the social cue. Although we found no strong evidence for such emotional salience transfer, our findings suggest that social information is flexibly incorporated into valence judgments based on the subjective uncertainty. This suggests that the influence of social cues in affective learning is metacognitively regulated, rather than automatically applied.

Several limitations should be acknowledged. First, our design relied on emotionally ambiguous stimuli, which may have introduced individual variability in how social cues were used or interpreted. This, in 46 turn, prompted exploratory analyses that used classification accuracy as a proxy for subjective uncertainty. Further, the experimental design complicates comparisons with prior studies using inherently neutral stimuli. Second, the use of emotional facial expressions as social cues—while ecologically valid—may have limited associative strength compared to more explicit reward or punishment outcomes. Third, although we preregistered our hypotheses and used equivalence testing to interpret null findings, our theoretical shift toward a metacognitive framework was developed post hoc. Thus, interpretations involving subjective uncertainty should be considered exploratory and require replication. Finally, ERP components like the EPN and LPC are influenced by both stimulus-driven and task-related factors, and disentangling these influences remains challenging in complex associative paradigms.

Taken together, our findings provide novel evidence that the influence of social cues on affective learning is shaped by metacognitive factors, particularly subjective uncertainty. Rather than transferring emotional salience automatically, social cues appear to act as flexible informational resources— modulating perceptual and memory processes depending on the availability and strength of internal evidence. These results extend current theories of social appraisal and emotional learning by highlighting the importance of individual-level uncertainty in guiding the integration of social information over time.

## Protocol Registration

The protocol was registered and is publicly available here: https://osf.io/tyq84

## Data availability

The behavioral raw data for the main experiment is available here https://osf.io/qrnxj/?view_only=baf5a10c98a44fa28c183782c5577acf. EEG data are not publicly available for privacy reasons (no consent from participants to publish the raw EEG data).

## Code availability

The code for the data simulation for power analysis, experiment script, pre-preprocessing and statistical analyses have been submitted https://osf.io/qrnxj/?view_only=baf5a10c98a44fa28c183782c5577ac.

## Acknowledgements

This work was supported by the RTG 2070, funded by DFG grant 254142454/GRK 2070 and CRC 1528 “Cognition of Interaction”. The funders have/had no role in study design, data collection and analysis, decision to publish or preparation of the manuscript. We thank Dr. Roger Mundry for his assistance with the power analysis and his recommendations on the analysis plans.

## Author contributions

SM – Conceptualization, Methodology, Writing – Original Draft

AS – Conceptualizing, Methodology, Writing – Original Draft, Supervision

## Competing interests

The authors declare no competing financial and non-financial interests.

## Notes

### Competing Interest Statement

The authors have declared no competing interest.

