## Supplement 1 for "Social learning of emotion and its implication for memory: An ERP Study"

### Supplementary Information 1

#### The timing for target image presentation to induce 'perceptual uncertainty'

The timing of the target image presentation – 27 ms – is chosen to create 'perceptual uncertainty' in judging the valence of the target images. Previously, it has been found that images presented for 20 ms are sufficient to detect the presence or absence of a particular target stimulus (animals) and that the visual processing of the target stimulus is reflected in the ERPs around 150 ms after stimulus onset [84]. A recent study using drift-diffusion modeling suggests that a stimulus presentation duration of 25 ms with backward masking achieves conscious perception, with a mean accuracy close to 80% in detecting the valence categories of facial expressions - happy, sad, or neutral [85]. Results from a visual awareness study indicated that at a stimulus presentation timing of 33 ms followed by backward masking, a considerable number of participants (64%) were able to detect the presence of an emotional target stimulus [86]. However, the above cited studies used a stimulus presence/absence detection task or a valence classification task in facial expressions – which is relatively easier as compared to valence classification of images.

In the present study, we aim to creating perceptual uncertainty about the valence of the target images while consciously perceiving the target images on the screen. Hence, we chose a stimulus presentation duration of 27 ms but with a forward mask to allow the visual processing of the target image even in its absence. The chosen timing of stimulus presentation presented with the forward mask, will enable conscious perception of the target image while creating perceptual uncertainty in judging the valence of the target images. Additionally, the chosen timing also fits well with the frame rate of the monitor that we will use for stimulus presentation (144 Hz frame rate, 27 ms equals 4 frames) to achieve a better stimulus presentation precision.

In our design, we propose to use a forward mask rather than a backward mask followed by a blank screen to allow the visual processing of the briefly presented target image to take place. Since we are interested in the neural dynamics of the target image at both the early and later stages of processing, having a forward mask would introduce considerable 'perceptual uncertainty' in judging the valence of the target images, while still allowing the processing of the target image to take place during the blank screen, which can be captured by the ERP measures time-locked to the target image presentation.

Taken together, the cumulative effect of the timing of the brief stimulus presentation timing along with the forward mask will make social cues such as facial expressions highly relevant for valence judgments of the target images.

#### Image selection criteria

| | Mean Valence | Mean Arousal | Mean Complexity |
|---|---|---|---|
| **Positive (N = 48)** | 6.96 (.03) | 4.72 (.06) | 2.10 (.07) |
| **Negative (N = 36)** | 2.55 (.08) | 5.67 (.08) | 2.29 (.06) |
| **Neutral (N = 41)** | 4.88 (.03) | 3.07 (.05) | 1.15 (.07) |

**Table S1.** Selected target stimuli with their mean Valence and Arousal ratings from the IAPS database (Lang et al., 2005), and mean complexity ratings from Bradley et al., (2007) study. Values in parentheses indicate standard errors.

#### Figures

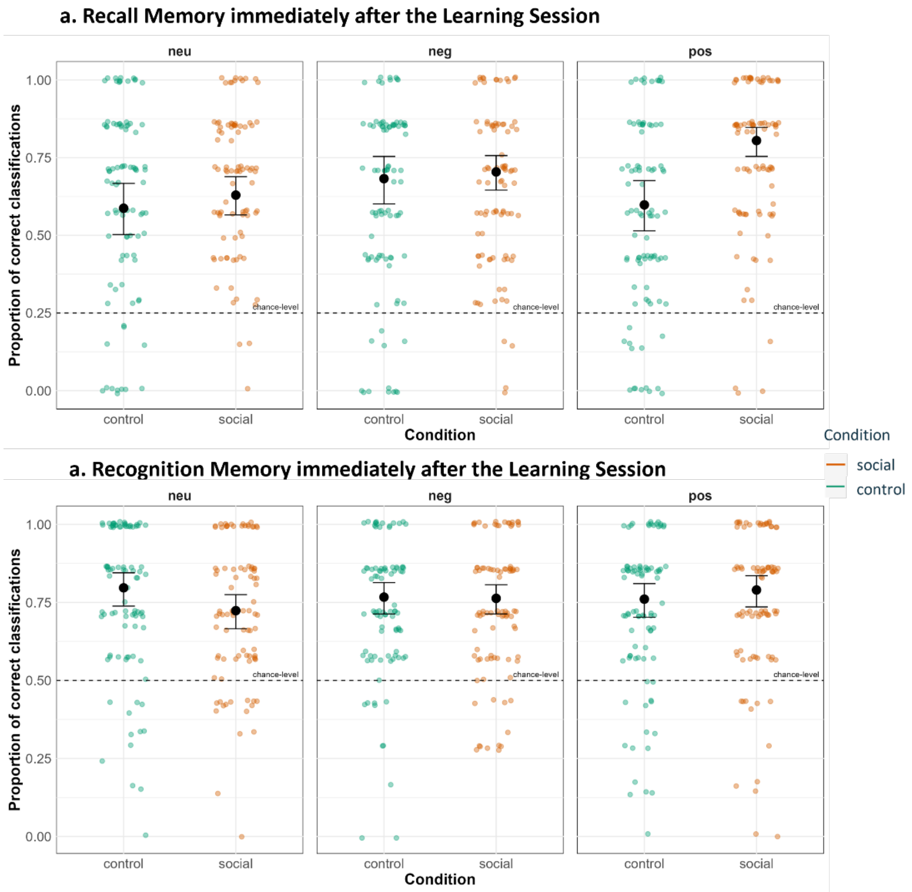


**Figure S1.** a. *Recall memory*. *b. Recognition memory.* Black dots indicate mean model-based predicted values along with the error bars indicating 95% CIs. Overlayed points represent mean accuracy values of individual participant per valence, per condition.

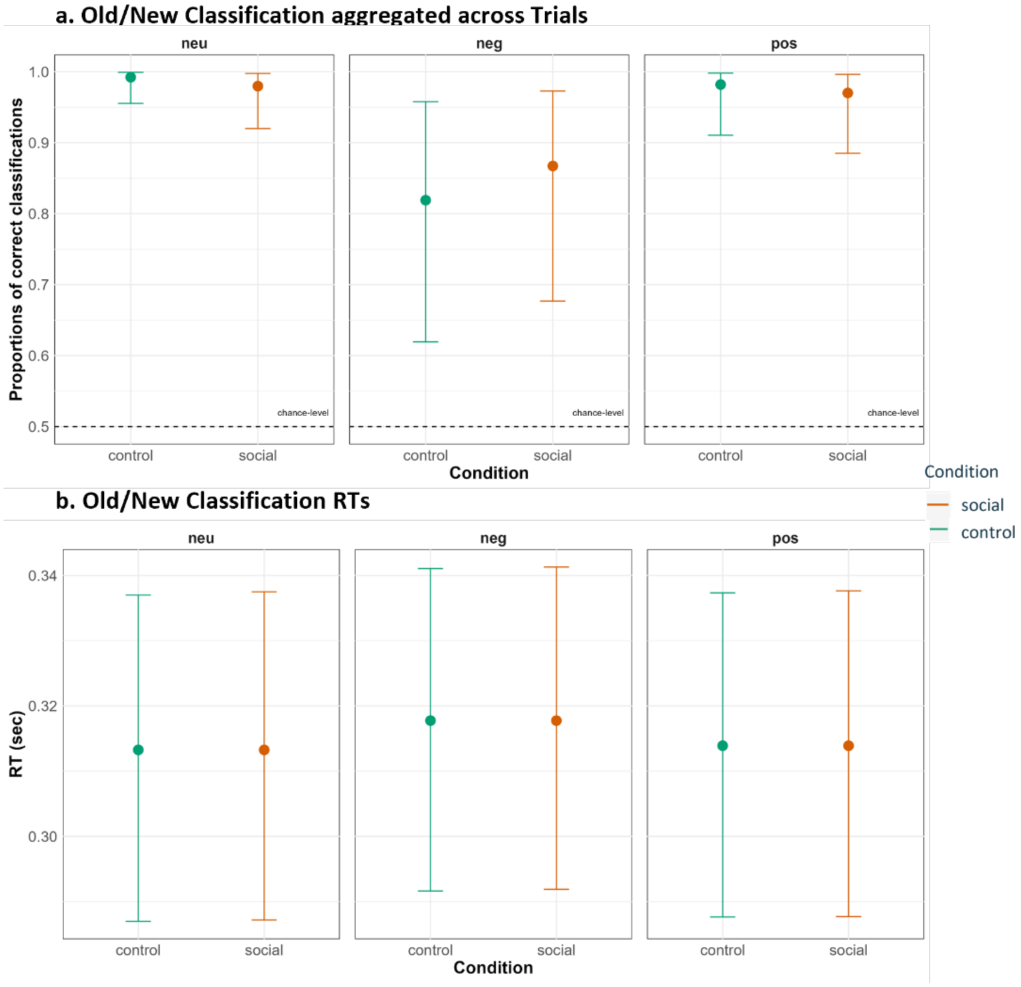


**Figure S2.** *a. Old/New Classification Accuracy aggregated across Trials. b. Old/New Classification RTs aggregated across Trials.* Plots represent mean model-based predicted values with error bars representing 95% CIs.

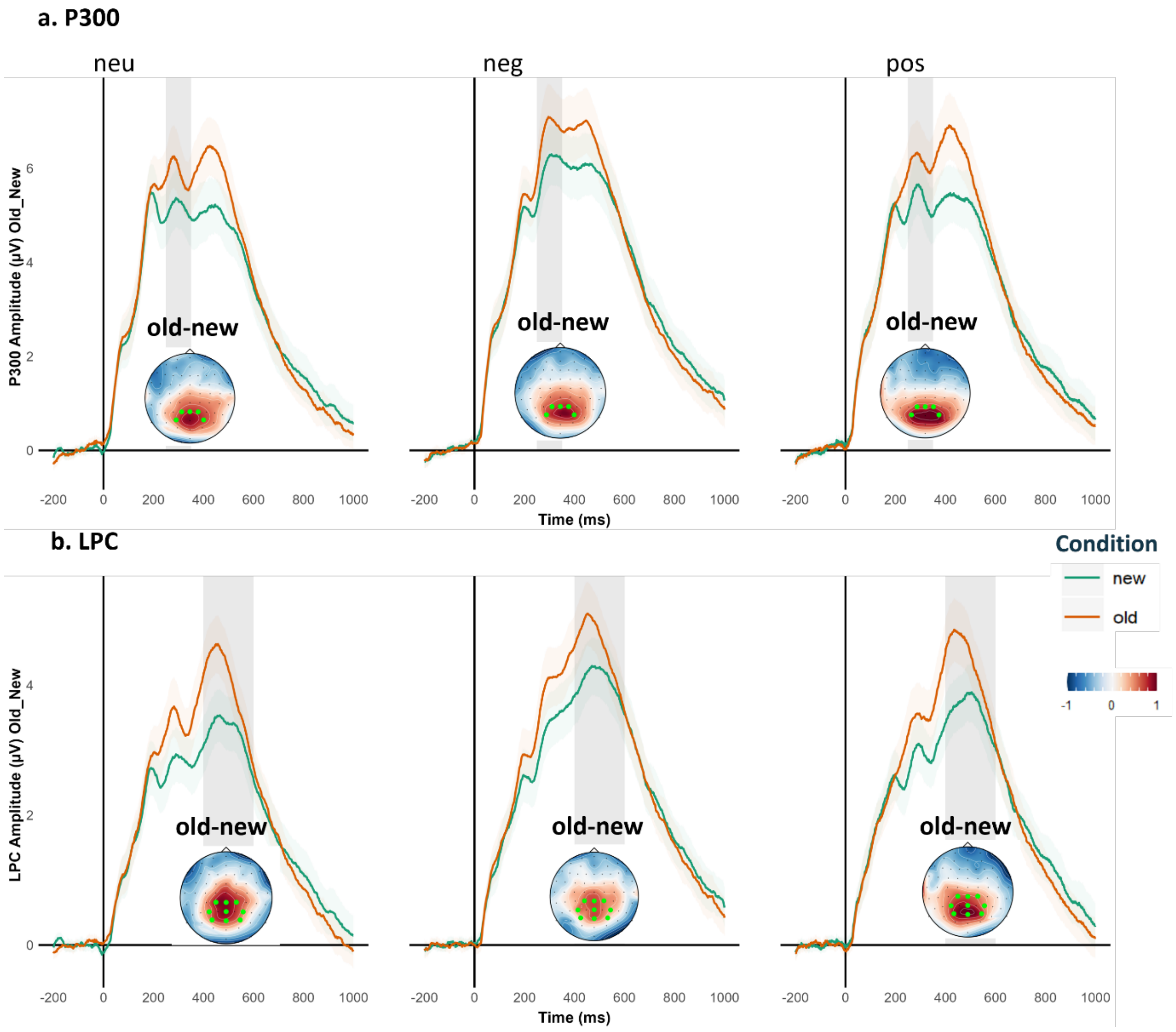


**Figure S3.** *a. P300. b. LPC*. Grand average waveforms across conditions for each of the target image valence category. Gray rectangular shaded region marks the time-window of the ERP. Topoplots indicate difference between conditions and green dots are the corresponding regions of interest (ROIs) for each of the ERPs.

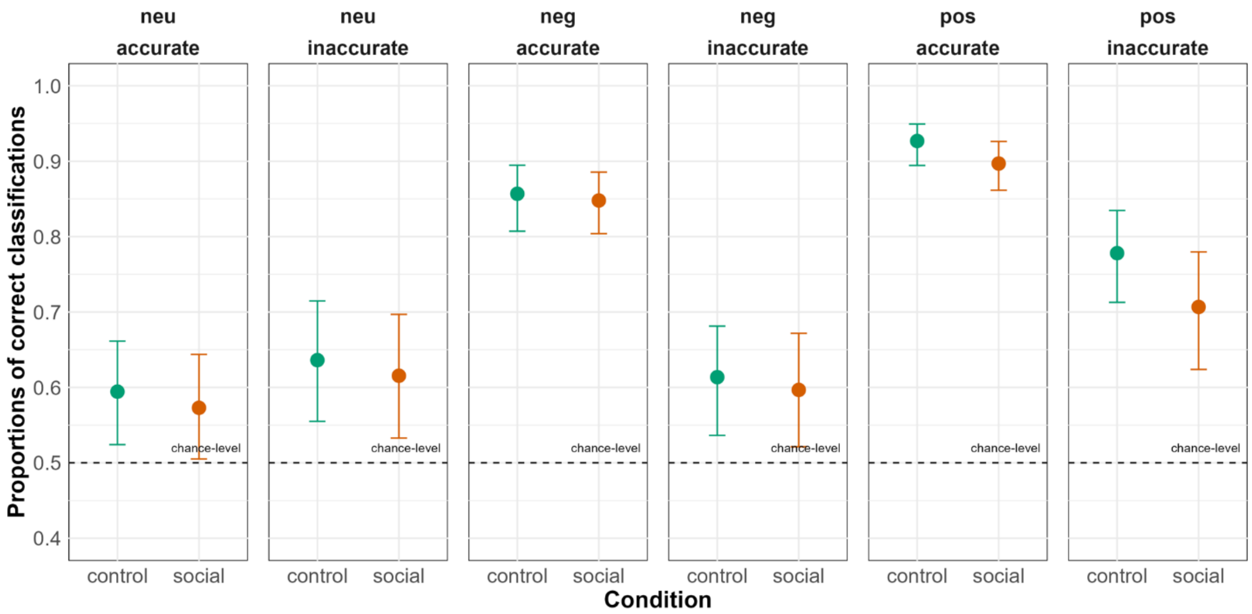


**Figure S4.** *Old-New Classification Accuracy split by classification accuracy*. Plots represent mean model-based predicted values with their corresponding 95% CIs as error bars.

#### Equivalence Testing

#### Day 1 Learning Session

**EPN:** To assess whether the non-significant condition effect could be considered practically equivalent to zero within a meaningful bound, we conducted a two one-sided tests (TOST) procedure using raw summary statistics. The equivalence test was significant, $t_{lower}(79) = 2.053$, $p = .022$; $\boldsymbol{t}_{upper}(79) = –27.273$, $p < .001$, indicating that the observed effect fell significantly within the equivalence bounds of –0.1 and +0.1. This supports the interpretation that the condition effect was small enough to be considered practically negligible.

**LPC:** To determine whether the non-significant condition effect could be considered practically negligible, we conducted a TOST procedure using raw summary statistics. The equivalence test was significant, $t_{lower}(79) = 24.150$, $p < .001$; $\boldsymbol{t}_{upper}(79) = –11.628$, $p < .001$, indicating that the observed effect fell significantly within the equivalence bounds of –0.1 and +0.1, suggesting that the effect was small enough to be considered practically equivalent to zero within the specified bounds.

#### Day 2 Test Session

**P1:** To evaluate whether this non-significant condition effect could be considered practically negligible, we conducted a TOST procedure using raw summary statistics. The equivalence test was significant, $t_{lower}(79)$

= 14.090, $p$ < .001; $\boldsymbol{t}_{\text{upper}}(79) = –10.415$, $p$ < .001, indicating that the observed effect fell significantly within the equivalence bounds of –0.1 and +0.1. This supports the conclusion that the condition effect was practically equivalent to zero within the specified bounds.

**LPC:** To evaluate whether the observed effect of condition was not only non-significant but also statistically equivalent to zero within a meaningful bound, we conducted a TOST procedure using raw summary statistics. The equivalence test was significant, $t_{\text{lower}}(79) = 28.085$, $p$ < .001; $\boldsymbol{t}_{\text{upper}}(79) = –7.692$, $p$ < .001, indicating that the observed effect fell significantly within the equivalence bounds of –0.1 and +0.1. This supports the conclusion that the condition effect was statistically equivalent to zero within the specified bounds.
