## Supplement 2 for "Social learning of emotion and its implication for memory: An ERP Study"

### Supplementary Information 2

### Results Tables

***Note:*** *p value = Satterthwaite approximated p values; CI = 95% confidence intervals; stab = model stability (estimate ranges leaving out one participant at a time – only done for exploratory analyses). Trials z-transformed to a mean of 0 and sd of 1; mean and sd of the original variable is 9.318 and 5.216 respectively. Image Complexity z-transformed to a mean of 0 and sd of 1; mean and sd of the original variable is 1.851 and 0.684 respectively. For all the pre-registered models, valence 'neutral' and condition 'control' are the reference levels. For the exploratory models, classification accuracy 'accurate' is the reference level.*

- *p-values of the intercepts not shown due to limited interpretability*
- *For the exploratory models, model stability was calculated using the DFBeta method by excluding levels of the grouping factor one at a time to see the effect on the model coefficients. We used the cut off ±2/sqrt(N), where N is the sample size to assess if the model estimates are unstable with respect to the specific predictor. Based on this, all of our model estimates were within the specified cut-off and therefore there was no model stability issue.*

##### Valence Classification Task (Day 1)

###### Table S1: Valence Classification Accuracy

| | estimate | std. error | z value | p value | lower CI | upper CI |
|---|---|---|---|---|---|---|
| (Intercept) | 2.266 | 0.436 | 5.199 | - | 1.432 | 3.100 |
| Valence negative | -3.371 | 0.644 | -5.233 | <.001 | -4.635 | -2.074 |
| Valence positive | -2.465 | 0.576 | -4.282 | <.001 | -3.678 | -1.358 |
| Condition social | 0.794 | 0.353 | 2.251 | .024 | 0.084 | 1.532 |
| Trials | 0.347 | 0.058 | 6.034 | <.001 | 0.227 | 0.456 |
| Image Complexity | -0.393 | 0.228 | -1.724 | .085 | -0.831 | 0.052 |
| Valence negative: Condition social | 1.667 | 0.425 | 3.919 | <.001 | 0.720 | 2.565 |
| Valence positive: Condition social | 1.551 | 0.421 | 3.684 | <.001 | 0.727 | 2.451 |
| Valence negative: Trials | -0.262 | 0.088 | -2.972 | .003 | -0.426 | -0.075 |
| Valence positive: Trials | -0.086 | 0.084 | -1.022 | .307 | -0.254 | 0.084 |

###### Table S2: Valence Classification RT

| | estimate | std. error | t value | p value | lower CI | upper CI |
|---|---|---|---|---|---|---|
| (Intercept) | 0.423 | 0.015 | 27.756 | - | 0.393 | 0.454 |
| Valence negative | -0.006 | 0.006 | -1.103 | .276 | -0.016 | 0.005 |
| Valence positive | -0.014 | 0.005 | -2.632 | .010 | -0.024 | -0.003 |
| Condition social | 0.004 | 0.007 | 0.603 | .550 | -0.010 | 0.018 |
| Trials | -0.045 | 0.005 | -9.593 | <.001 | -0.055 | -0.036 |
| Image Complexity | 0.008 | 0.002 | 3.836 | <.001 | 0.004 | 0.012 |
| Valence negative: Condition social | 0.008 | 0.006 | 1.448 | .156 | -0.002 | 0.020 |

| | | | | | | |
|---|---|---|---|---|---|---|
| Valence positive: Condition social | 0.012 | 0.006 | 2.111 | .040 | 0.001 | 0.023 |

**Table S3: P1 ERP**

| | estimate | std. error | t value | p value | lower CI | upper CI |
|---|---|---|---|---|---|---|
| (Intercept) | 7.268 | 0.362 | 20.075 | - | 6.554 | 8.012 |
| Valence negative | -0.069 | 0.159 | -0.435 | .669 | -0.347 | 0.231 |
| Valence positive | -0.027 | 0.148 | -0.181 | .858 | -0.306 | 0.268 |
| Condition social | -0.118 | 0.065 | -1.804 | .079 | -0.250 | 0.009 |
| Trials | 0.136 | 0.057 | 2.389 | .020 | 0.029 | 0.255 |
| Image Complexity | 0.006 | 0.066 | 0.090 | .930 | -0.118 | 0.136 |

**Table S4: EPN ERP**

| | estimate | std. error | t value | p value | lower CI | upper CI |
|---|---|---|---|---|---|---|
| (Intercept) | -0.422 | 0.353 | -1.196 | - | -1.143 | 0.271 |
| Valence negative | -0.779 | 0.369 | -2.114 | .039 | -1.567 | -0.069 |
| Valence positive | -0.595 | 0.328 | -1.813 | .076 | -1.260 | 0.053 |
| Trials | 0.133 | 0.061 | 2.161 | .034 | 0.011 | 0.252 |
| Condition social | -0.086 | 0.061 | -1.409 | .166 | -0.202 | 0.042 |
| Image Complexity | 0.943 | 0.155 | 6.080 | .000 | 0.639 | 1.272 |
| Valence negative: Trials | -0.146 | 0.056 | -2.585 | .012 | -0.260 | -0.031 |
| Valence positive: Trials | -0.141 | 0.062 | -2.255 | .028 | -0.262 | -0.021 |

**Table S5: LPC ERP**

| | estimate | std. error | t value | p value | lower CI | upper CI |
|---|---|---|---|---|---|---|
| (Intercept) | 1.348 | 0.254 | 5.315 | - | 0.859 | 1.814 |
| Valence negative | 0.359 | 0.212 | 1.687 | .098 | -0.057 | 0.771 |
| Valence positive | 0.555 | 0.191 | 2.905 | .005 | 0.183 | 0.927 |
| Trials | 0.136 | 0.053 | 2.588 | .012 | 0.032 | 0.239 |
| Condition social | 0.035 | 0.050 | 0.697 | .490 | -0.062 | 0.136 |
| Image Complexity | -0.272 | 0.090 | -3.014 | .003 | -0.444 | -0.097 |

#### Old/New Classification Task (Day 2)

##### Table S6 : Old/New Classification Accuracy

| | estimate | std. error | z value | p value | lower CI | upper CI |
|---|---|---|---|---|---|---|
| (Intercept) | 3.915 | 1.133 | 3.456 | - | 2.500 | 9.763 |
| Valence negative | 1.898 | 1.636 | 1.160 | .246 | -2.137 | 5.469 |
| Valence positive | 5.354 | 1.461 | 3.666 | <.001 | 1.053 | 9.061 |
| Condition social | -1.154 | 0.530 | -2.179 | .029 | -3.022 | 0.014 |
| Trials | 0.551 | 0.182 | 3.021 | .003 | 0.165 | 0.769 |
| Image Complexity | -4.300 | 0.656 | -6.559 | .000 | -6.476 | -1.031 |
| Valence negative: Condition social | 1.657 | 0.735 | 2.254 | .024 | 0.046 | 5.381 |
| Valence positive: Condition social | 0.270 | 0.744 | 0.363 | .717 | -1.704 | 2.205 |
| Valence negative: Trials | -0.292 | 0.244 | -1.196 | .232 | -0.676 | 0.190 |
| Valence positive: Trials | -0.158 | 0.248 | -0.638 | .524 | -0.524 | 0.255 |
| Condition social: Trials | -0.163 | 0.215 | -0.757 | .449 | -0.469 | 0.242 |
| Valence negative: Condition social: Trials | 0.274 | 0.300 | 0.914 | .361 | -0.306 | 0.713 |
| Valence positive: Condition social: Trials | 0.745 | 0.301 | 2.479 | .013 | 0.196 | 1.156 |

##### Table S7 : Old/New Response Time

| | estimate | std. error | t value | p value | lower CI | upper CI |
|---|---|---|---|---|---|---|
| (Intercept) | 0.313 | 0.012 | 25.272 | - | 0.288 | 0.336 |
| Valence negative | 0.004 | 0.003 | 1.723 | .092 | -0.001 | 0.010 |
| Valence positive | 0.001 | 0.003 | 0.226 | .824 | -0.005 | 0.006 |
| Condition social | 0.000 | 0.001 | -0.003 | .999 | -0.003 | 0.003 |
| Trials | -0.048 | 0.004 | -12.750 | <.001 | -0.054 | -0.040 |
| Image Complexity | 0.004 | 0.001 | 3.347 | .001 | 0.002 | 0.006 |

**Table S8: P1 ERP**

| | estimate | std. error | t value | p value | lower CI | upper CI |
|---|---|---|---|---|---|---|
| (Intercept) | 11.311 | 0.627 | 18.050 | - | 10.126 | 12.526 |
| Valence negative | 0.345 | 0.240 | 1.439 | .134 | -0.138 | 0.840 |
| Valence positive | 0.068 | 0.222 | 0.306 | .726 | -0.367 | 0.502 |
| Condition social | 0.015 | 0.073 | 0.211 | .746 | -0.120 | 0.157 |
| Trials | 0.365 | 0.071 | 5.152 | <.001 | 0.232 | 0.506 |
| Image Complexity | 0.024 | 0.101 | 0.233 | .833 | -0.184 | 0.229 |

**Table S9: EPN ERP**

| | estimate | std. error | t value | p value | lower CI | upper CI |
|---|---|---|---|---|---|---|
| (Intercept) | 10.545 | 0.579 | 18.215 | - | 9.435 | 11.703 |
| Valence negative | -0.800 | 0.455 | -1.759 | .085 | -1.710 | 0.031 |
| Valence positive | -0.398 | 0.412 | -0.966 | .342 | -1.259 | 0.393 |
| Trials | 0.878 | 0.082 | 10.696 | <.001 | 0.732 | 1.031 |
| Condition social | 0.312 | 0.121 | 2.575 | .013 | 0.079 | 0.532 |
| Image Complexity | 0.620 | 0.187 | 3.325 | .001 | 0.245 | 0.982 |
| Valence negative: Trials | -0.148 | 0.072 | -2.070 | .044 | -0.297 | -0.014 |
| Valence positive: Trials | -0.151 | 0.069 | -2.195 | .033 | -0.290 | -0.030 |
| Valence negative: Condition social | -0.247 | 0.172 | -1.438 | .163 | -0.562 | 0.087 |
| Valence positive: Condition social | -0.422 | 0.172 | -2.453 | .017 | -0.763 | -0.109 |

**Table S10: LPC ERP**

| | estimate | std. error | t value | p value | lower CI | upper CI |
|---|---|---|---|---|---|---|
| (Intercept) | 3.943 | 0.235 | 16.771 | - | 3.480 | 4.408 |
| Valence negative | 0.652 | 0.164 | 3.967 | <.001 | 0.332 | 0.996 |
| Valence positive | 0.360 | 0.148 | 2.431 | .018 | 0.068 | 0.643 |
| Trials | -0.132 | 0.066 | -1.995 | .050 | -0.258 | -0.009 |
| Condition social | 0.057 | 0.050 | 1.126 | .263 | -0.045 | 0.151 |
| Image Complexity | -0.095 | 0.069 | -1.390 | .172 | -0.232 | 0.041 |
| Valence negative: Trials | -0.132 | 0.062 | -2.111 | .040 | -0.256 | -0.004 |
| Valence positive: Trials | -0.051 | 0.054 | -0.945 | .350 | -0.160 | 0.060 |

### Exploratory Analyses

#### Valence Classification Task (Day 1)

##### Table S11: RTs accounting for classification accuracy

| | estimate | std. error | t value | p value | stab min | stab max | lower CI | upper CI |
|---|---|---|---|---|---|---|---|---|
| (Intercept) | 0.427 | 0.017 | 24.475 | - | 0.422 | 0.430 | 0.393 | 0.462 |
| Valence negative | -0.017 | 0.012 | -1.447 | .150 | -0.022 | -0.013 | -0.039 | 0.006 |
| Valence positive | -0.024 | 0.012 | -2.076 | .040 | -0.027 | -0.021 | -0.047 | 0.000 |
| Condition social | 0.017 | 0.010 | 1.663 | .100 | 0.012 | 0.020 | -0.001 | 0.038 |
| Class. accuracy inaccurate | 0.037 | 0.012 | 3.003 | .003 | 0.033 | 0.041 | 0.012 | 0.060 |
| Image Complexity | 0.006 | 0.004 | 1.561 | .123 | 0.004 | 0.006 | -0.002 | 0.012 |
| Valence negative: Condition social | 0.013 | 0.011 | 1.202 | .228 | 0.010 | 0.016 | -0.009 | 0.034 |
| Valence positive: Condition social | 0.016 | 0.012 | 1.281 | .205 | 0.010 | 0.021 | -0.008 | 0.040 |
| Valence negative: Class. accuracy inaccurate | 0.021 | 0.015 | 1.416 | .163 | 0.016 | 0.029 | -0.008 | 0.049 |
| Valence positive: Class. accuracy inaccurate | 0.057 | 0.014 | 4.041 | <.001 | 0.052 | 0.060 | 0.031 | 0.083 |

##### Table S12: P1 Amplitudes Accounting for Classification Accuracy

| | estimate | std. error | t value | p value | stab min | stab max | lower CI | upper CI |
|---|---|---|---|---|---|---|---|---|
| (Intercept) | 5.367 | 0.362 | 14.845 | - | 5.262 | 5.459 | 4.664 | 6.041 |
| Valence negative | 0.476 | 0.258 | 1.845 | .066 | 0.344 | 0.519 | -0.047 | 0.970 |
| Valence positive | 0.098 | 0.245 | 0.399 | .692 | -0.011 | 0.147 | -0.396 | 0.540 |
| Condition social | -0.266 | 0.211 | -1.261 | .208 | -0.318 | -0.212 | -0.704 | 0.169 |
| Class. accuracy inaccurate | 0.763 | 0.249 | 3.064 | .002 | 0.666 | 0.827 | 0.278 | 1.244 |
| Image Complexity | 0.066 | 0.068 | 0.970 | .331 | 0.048 | 0.087 | -0.060 | 0.203 |
| Valence negative: Condition social | -0.409 | 0.308 | -1.328 | .185 | -0.468 | -0.323 | -1.021 | 0.159 |
| Valence positive: Condition social | -0.044 | 0.308 | -0.142 | .887 | -0.106 | 0.071 | -0.659 | 0.593 |
| Valence negative: Class. accuracy inaccurate | -1.621 | 0.351 | -4.617 | <.001 | -1.698 | -1.419 | -2.278 | -0.902 |
| Valence positive: Class. accuracy inaccurate | -0.988 | 0.343 | -2.885 | .004 | -1.092 | -0.857 | -1.622 | -0.287 |
| Condition social: Class. accuracy inaccurate | -0.229 | 0.348 | -0.657 | .512 | -0.289 | -0.148 | -0.904 | 0.438 |
| Valence negative: Condition social: Class. accuracy inaccurate | 1.280 | 0.468 | 2.737 | .006 | 1.049 | 1.391 | 0.407 | 2.215 |
| Valence positive: Condition social: Class. accuracy inaccurate | 0.900 | 0.479 | 1.877 | .060 | 0.694 | 1.070 | -0.084 | 1.833 |

##### Table S13: EPN Amplitudes Accounting for Classification Accuracy

| | estimate | std. error | t value | p value | stab min | stab max | lower CI | upper CI |
|---|---|---|---|---|---|---|---|---|
| (Intercept) | -0.393 | 0.273 | -1.440 | - | -0.465 | -0.332 | -0.896 | 0.148 |
| Valence negative | -0.778 | 0.146 | -5.318 | <.001 | -0.813 | -0.740 | -1.060 | -0.477 |
| Valence positive | -0.582 | 0.145 | -4.023 | <.001 | -0.636 | -0.542 | -0.848 | -0.292 |
| Class. accuracy inaccurate | 0.146 | 0.094 | 1.549 | .131 | 0.112 | 0.170 | -0.039 | 0.340 |
| Condition social | -0.180 | 0.098 | -1.840 | .069 | -0.210 | -0.149 | -0.360 | 0.004 |
| Image Complexity | 0.889 | 0.070 | 12.679 | <.001 | 0.866 | 0.908 | 0.755 | 1.031 |

**Table S14: LPC Amplitudes Accounting for Classification Accuracy**

| | estimate | std. error | t value | p value | stab min | stab max | lower CI | upper CI |
|---|---|---|---|---|---|---|---|---|
| (Intercept) | 1.178 | 0.228 | 5.175 | - | 1.111 | 1.247 | 0.745 | 1.643 |
| Valence negative | 0.678 | 0.157 | 4.313 | <.001 | 0.626 | 0.728 | 0.367 | 0.989 |
| Valence positive | 0.963 | 0.154 | 6.268 | <.001 | 0.920 | 0.999 | 0.656 | 1.259 |
| Class. accuracy inaccurate | -0.235 | 0.153 | -1.536 | .128 | -0.286 | -0.160 | -0.542 | 0.087 |
| Condition social | -0.066 | 0.076 | -0.874 | .389 | -0.099 | -0.049 | -0.211 | 0.083 |
| Image Complexity | -0.228 | 0.055 | -4.178 | <.001 | -0.251 | -0.205 | -0.334 | -0.122 |
| Valence negative: Class. accuracy inaccurate | -0.582 | 0.205 | -2.838 | .006 | -0.645 | -0.482 | -0.967 | -0.193 |
| Valence positive: Class. accuracy inaccurate | -1.131 | 0.226 | -4.996 | <.001 | -1.201 | -1.054 | -1.590 | -0.699 |

#### Old/New Classification Task (Day 2)

**Table S15: Old/New Classification Accuracy**

| | estimate | std. error | z value | p value | stab min | stab max | lower CI | upper CI |
|---|---|---|---|---|---|---|---|---|
| (Intercept) | 0.382 | 0.148 | 2.580 | - | 0.344 | 0.434 | 0.097 | 0.670 |
| Valence negative | 1.407 | 0.229 | 6.135 | <.001 | 1.349 | 1.487 | 0.978 | 1.854 |
| Valence positive | 2.157 | 0.250 | 8.613 | <.001 | 2.068 | 2.234 | 1.706 | 2.645 |
| Condition social | -0.088 | 0.146 | -0.605 | .545 | -0.117 | -0.035 | -0.360 | 0.190 |
| Class. accuracy inaccurate | 0.177 | 0.167 | 1.057 | .290 | 0.119 | 0.224 | -0.163 | 0.491 |
| Image Complexity | -0.705 | 0.070 | -10.119 | <.001 | -0.728 | -0.685 | -0.850 | -0.576 |
| Valence negative: Condition social | 0.017 | 0.213 | 0.082 | .935 | -0.076 | 0.056 | -0.407 | 0.420 |
| Valence positive: Condition social | -0.287 | 0.253 | -1.131 | .258 | -0.370 | -0.230 | -0.777 | 0.196 |
| Valence negative: Class. accuracy inaccurate | -1.503 | 0.240 | -6.276 | <.001 | -1.570 | -1.415 | -1.969 | -1.051 |
| Valence positive: Class. accuracy inaccurate | -1.461 | 0.255 | -5.734 | <.001 | -1.574 | -1.373 | -1.981 | -1.006 |

#### Table S16: EPN Amplitudes Accounting for Classification Accuracy

| | estimate | std. error | t value | p value | stab min | stab max | lower CI | upper CI |
|---|---|---|---|---|---|---|---|---|
| (Intercept) | 10.594 | 0.514 | 20.626 | - | 10.428 | 10.743 | 9.628 | 11.743 |
| Valence negative | -1.534 | 0.265 | -5.797 | <.001 | -1.649 | -1.402 | -2.030 | -1.044 |
| Valence positive | -0.367 | 0.235 | -1.564 | .119 | -0.428 | -0.285 | -0.821 | 0.087 |
| Condition social | 0.088 | 0.179 | 0.490 | .625 | 0.043 | 0.122 | -0.273 | 0.439 |
| Class. accuracy inaccurate | -0.031 | 0.234 | -0.134 | .887 | -0.112 | 0.054 | -0.476 | 0.443 |
| Image Complexity | 0.677 | 0.065 | 10.469 | <.001 | 0.659 | 0.697 | 0.546 | 0.812 |
| Valence negative: Condition social | 0.391 | 0.317 | 1.235 | .222 | 0.297 | 0.574 | -0.221 | 1.038 |
| Valence positive: Condition social | -0.526 | 0.277 | -1.900 | .058 | -0.607 | -0.442 | -1.067 | 0.051 |
| Valence negative: Class. accuracy inaccurate | 0.765 | 0.351 | 2.179 | .031 | 0.595 | 0.879 | 0.075 | 1.411 |
| Valence positive: Class. accuracy inaccurate | -0.076 | 0.314 | -0.242 | .816 | -0.172 | 0.042 | -0.730 | 0.586 |
| Condition social: Class. accuracy inaccurate | 0.479 | 0.322 | 1.488 | .138 | 0.375 | 0.539 | -0.148 | 1.084 |
| Valence negative: Condition social: Class. accuracy inaccurate | -0.978 | 0.470 | -2.082 | .040 | -1.143 | -0.782 | -1.846 | -0.099 |
| Valence positive: Condition social: Class. accuracy inaccurate | -0.029 | 0.441 | -0.066 | .946 | -0.176 | 0.094 | -0.848 | 0.824 |

#### Table S17: LPC Amplitudes Accounting for Classification Accuracy

| | estimate | std. error | t value | p value | stab min | stab max | lower CI | upper CI |
|---|---|---|---|---|---|---|---|---|
| (Intercept) | 3.809 | 0.225 | 16.928 | - | 3.745 | 3.859 | 3.389 | 4.264 |
| Valence negative | 1.203 | 0.141 | 8.502 | <.001 | 1.154 | 1.245 | 0.904 | 1.471 |
| Valence positive | 0.689 | 0.132 | 5.213 | <.001 | 0.642 | 0.719 | 0.416 | 0.949 |
| Class. accuracy inaccurate | 0.296 | 0.107 | 2.776 | .006 | 0.274 | 0.321 | 0.084 | 0.497 |
| Condition social | 0.149 | 0.093 | 1.605 | .110 | 0.115 | 0.170 | -0.028 | 0.328 |
| Image Complexity | -0.077 | 0.041 | -1.867 | .063 | -0.088 | -0.059 | -0.159 | 0.004 |
| Valence negative: Class. accuracy inaccurate | -0.830 | 0.145 | -5.723 | <.001 | -0.869 | -0.799 | -1.095 | -0.534 |
| Valence positive: Class. accuracy inaccurate | -0.637 | 0.144 | -4.429 | <.001 | -0.681 | -0.603 | -0.892 | -0.356 |
| Valence negative: Condition social | -0.272 | 0.132 | -2.056 | .043 | -0.308 | -0.212 | -0.539 | 0.001 |
| Valence positive: Condition social | -0.171 | 0.140 | -1.225 | .225 | -0.218 | -0.128 | -0.437 | 0.115 |
